# Assignment of virus and antimicrobial resistance genes to microbial hosts in a complex microbial community by combined long-read assembly and proximity ligation

**DOI:** 10.1101/491175

**Authors:** Derek M. Bickhart, Mick Watson, Sergey Koren, Kevin Panke-Buisse, Laura M. Cersosimo, Maximilian O. Press, Curtis P. Van Tassell, Jo Ann S. Van Kessel, Bradd J. Haley, Seon Woo Kim, Cheryl Heiner, Garret Suen, Kiranmayee Bakshy, Ivan Liachko, Shawn T. Sullivan, Jay Ghurye, Mihai Pop, Paul J. Weimer, Adam M. Phillippy, Timothy P.L. Smith

## Abstract

The characterization of microbial communities by metagenomic approaches has been enhanced by recent improvements in short-read sequencing efficiency and assembly algorithms. We describe the results of adding long-read sequencing to the mix of technologies used to assemble a highly complex cattle rumen microbial community, and compare the assembly to current short read-based methods applied to the same sample. Contigs in the long-read assembly were 7-fold longer on average, and contained 7-fold more complete open reading frames (ORF), than the short read assembly, despite having three-fold lower sequence depth. The linkages between long-read contigs, provided by proximity ligation data, supported identification of 188 novel viral-host associations in the rumen microbial community that suggest cross-species infectivity of specific viral strains. The improved contiguity of the long-read assembly also identified 94 antimicrobial resistance genes, compared to only seven alleles identified in the short-read assembly. Overall, we demonstrate a combination of experimental and computational methods that work synergistically to improve characterization of biological features in a highly complex rumen microbial community.

## Background

Microbial genome assembly from metagenomic sequence of complex communities produces large numbers of genome fragments, rather than complete circular genomes, despite continuous improvements in methodology (1,2). Assembly is complicated by sequences that may occur repeatedly within strains (“repeats”) or shared among similar strains of bacterial and archaeal species, creating “branches” in the assembly graph that precludes accurate representation of individual component genomes, particularly when multiple closely-related strains of a species are present in the environment (3). Repetitive content contributes to difficulty in multicellular Eukaryotic genome assembly as well (4), but the problem becomes more complicated in metagenome assembly (5) due to the wide range of abundance among bacterial species and strains, and the presence of other environmental DNA (e.g. plants, protists).

The application of long-read sequencing appears to be a potential solution to many of the difficulties inherent to metagenomic assembly. Read lengths that exceed the size of highly repetitive sequences, such as ribosomal RNA gene clusters, have been shown to improve contig lengths in the initial assembly (6,7). However, longer repetitive regions are only capable of being completely resolved by long reads of equal or greater size to the repeat, which makes input DNA quality a priority in sequence library construction. This can present a problem in metagenomics samples as material-adherent bacterial populations produce tough extracellular capsules that require vigorous mechanical stress for lysis, resulting in substantial DNA fragmentation and single-strand nicks (8). Long-read sequencing technologies have been previously used in the assembly of the skin microbiome (9), several environmental metagenomes (10), and in the binning of contigs from a biogas reactor12; however, each of these projects has relied on additional coverage from short-read data to compensate for lower long-read coverage. Additionally, higher depths of coverage of long-reads from current generation sequencing technologies are necessary to overcome high, relative error rates that can impact assembly quality and influence functional genomic annotation (11). Still, there is substantial interest in generating assemblies derived from longer reads to enable better characterization of environmental and complex metagenomics communities (10). Metagenome WGS assemblies consisting entirely of long-reads have yet to be fully characterized, particularly those from complex, multi-kingdom symbiotic communities.

The bovine rumen is an organ that serves as the site of symbiosis between the cow and microbial species from all three taxonomic Superkingdoms of life that are dedicated to the degradation of highly recalcitrant plant polymers (12). With efficiency unrivaled by most abiotic industrial processes, the protists, archaea, bacteria and fungi that make up the rumen microbial community are able to process cellulose and other plant biopolymers into byproducts, such as volatile fatty acids (VFA), that can be utilized by the host. This process is supplemented by relatively minimal energy inputs, such as the basal body temperature of the host cow and the energy-efficient mastication of digesting plant material. The presence of organisms from all major Superkingdoms in varying degrees of abundance makes the rumen an excellent model for a complex, partially-characterized metagenome system. Assessments of rumen microbial presence and abundance have generally been limited to 16S rRNA amplicon sequencing (13–15); however, recent genome assemblies of metagenomic samples (16,17) or isolates (18) derived from the rumen provide suitable standards for the comparison of new assembly methods and techniques.

In this study, we compare and contrast several different technologies that are suitable for metagenome assembly and binning, and we highlight distinct biological features that each technology is able to best resolve. We show that contigs generated using longer-read sequencing tend to be larger than those generated by shorter-read sequencing methods; long-reads assemble more full-length genes and antimicrobial resistance gene alleles; and that long-reads can be suitable for identifying the host-specificity of assembled viruses/prophages in a metagenomics community. We also highlight novel host-viral associations and the potential horizontal transfer of antimicrobial resistance genes (ARG) in rumen microbial species using a combination of long-reads and Hi-C intercontig link data. Our data suggests that future metagenomics surveys should include a combination of different sequencing and conformational capture technologies in order to fully assess the diversity and biological functionality of a sample.

## Results

### Sample extraction quality and *de novo* genome assemblies

We extracted high molecular weight DNA from a combined rumen fluid and solid sample taken from a single, multiparous, cannulated cow and sequenced that sample using a short-read and a long-read DNA sequencing technology (see methods; Fig 1a). The short-read and long-read data were assembled separately and generated de novo assemblies with contig N100K counts (the number of contigs with lengths greater than 100 kbp) of 88 and 384, respectively (Table 1). While the long-read assembly was mostly comprised of larger contigs, the short-read assembly contained five-fold more assembled bases (1.0 gigabases vs. 5.1 gigabases). We also observed a slight bias in the GC content of assembled contigs, with the short-read assembly having a larger sampling of different, average GC content tranches than the long-read assembly in observed, assembled contigs (Fig 1b). Interestingly, the average GC content of the error-corrected long-reads indicated a bimodal distribution at the 0.5 and 0.25 ratios (Figure 1b) that is less pronounced in the GC statistics of the raw short-reads and both sets of assembly contigs. There are several possibilities for this discrepancy; however, it is possible that this lower GC content range belongs to unassembled protist or anaerobic fungi genomes which are known to be highly repetitive, and have low GC content (19,20).

**Figure 1.**
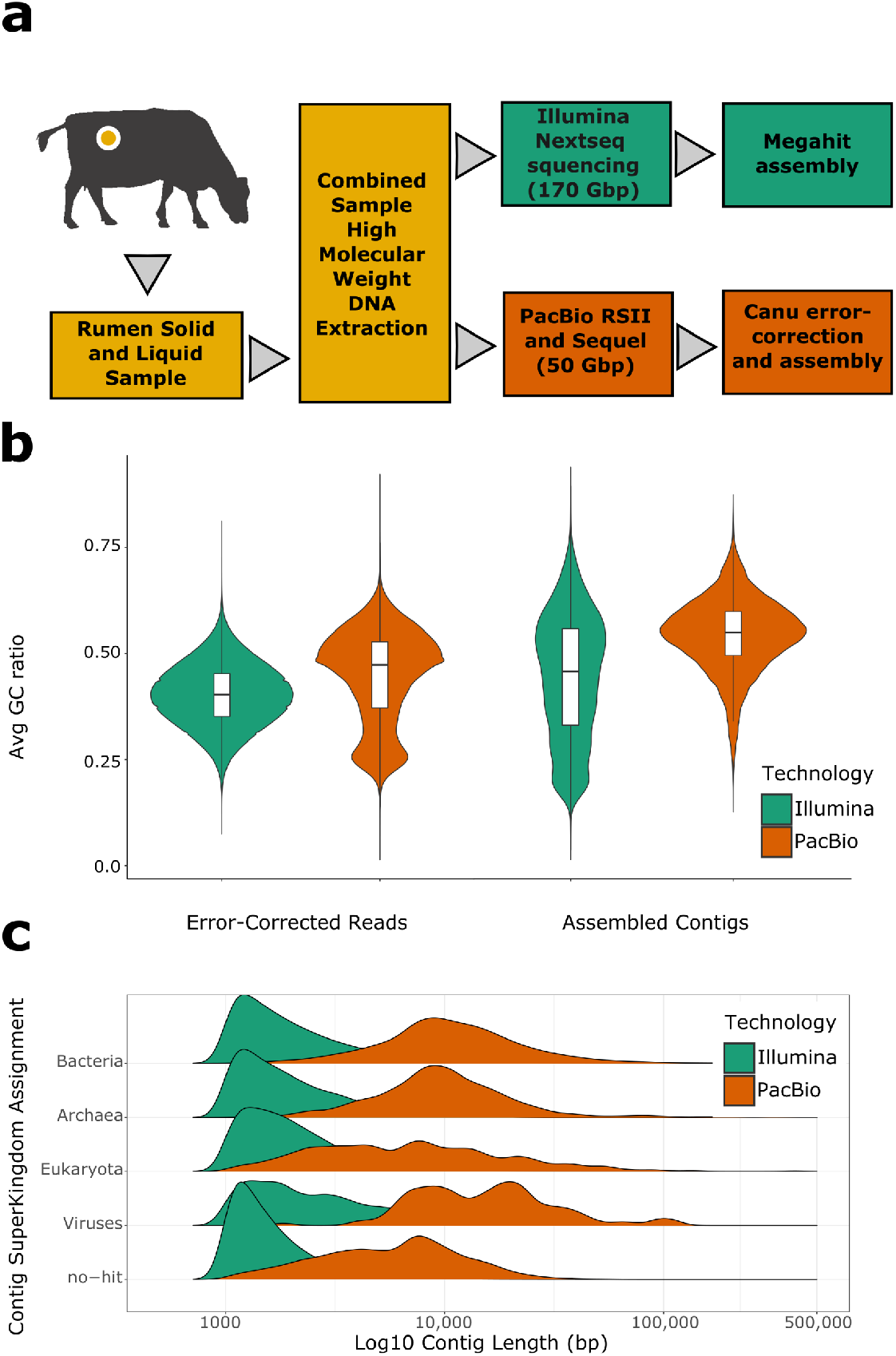
Assembly workflow and sampling bias estimates show GC% discrepancies in long-reads vs assemblies. Using the same sample from a cannulated cow, (A) we extracted DNA using a modified bead beating protocol that still preserved a large proportion of high molecular weight DNA strands. This DNA extraction was sequenced on a short-read sequencer (Illumina; dark green) and a long-read sequencer (PacBio RSII and Sequel; dark orange), with each sequence source assembled separately. Assessments of read- and contig-level GC% bias (B) revealed that a substantial proportion of sampled low GC DNA was not incorporated into either assembly. (C) Assembly contigs were annotated for likely superkingdoms of origin and were compared for overall contig lengths. The long-read assembly tended to have longer average contigs for each assembled superkingdom compared to the short-read assembly.

**Table 1.**
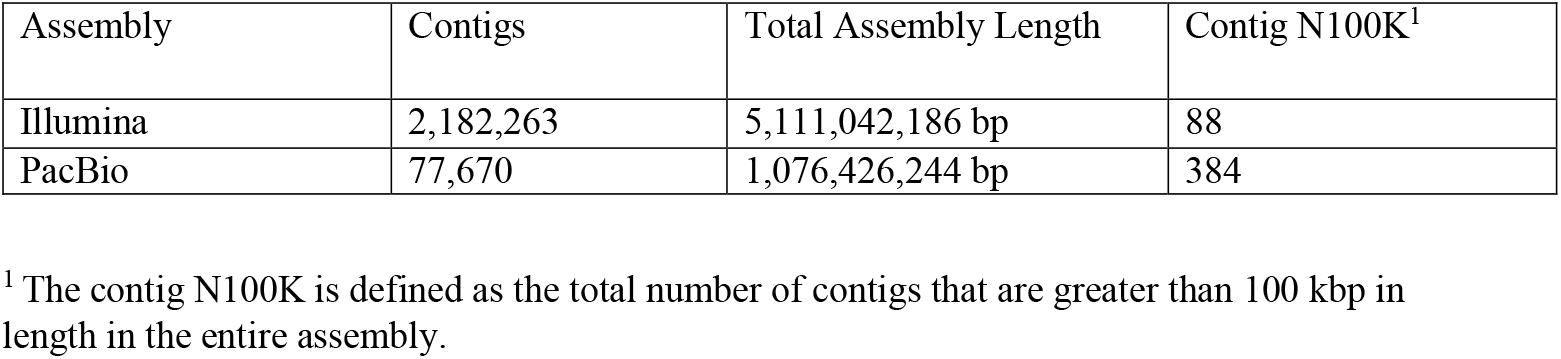
Assembly statistics

We noticed a slight discrepancy in the Superkingdom-specific contig lengths that suggests that many of our contigs of potential Eukaryotic-origins are shorter than those of the Bacteria and Archaea, whichcoincided with our observation of GC content bias in the assembly (Fig 1c). To assess the bias in GC content in our assembly of the long-read data, we calculated the overlap of raw long-reads with our long-read assembly contigs. Density estimates of long-reads that were not included in the long-read assembly (zero overlaps) mirrored the bimodal distribution of GC content in the raw long-reads previously observed, suggesting that a larger proportion of lower GC content reads had insufficient coverage to be assembled (Additional file 1: Fig S1). Furthermore, we note that the error corrected long-reads were filtered based on intra-dataset overlaps, resulting in a further reduction of bases compared with the starting, raw long-reads. The correction step removed 10% of the total reads for being singleton observations (zero overlaps with any other read) and trimmed the ends of 26% of the reads for having less than 2 overlaps. This may have also impacted the assembly of low abundance or highly complex genomes in the sample by removing rare observations of DNA sequence. We attempted to combine both the long-read and short-read datasets into a hybrid assembly; however, all attempts using currently available software were unsuccessful as currently available tools had prohibitive memory or runtime requirements due to the size of our input assemblies. We also investigated the use of long-reads in multiple-datasource scaffolding programs and found only minor improvements in assembly size that were achieved through the inclusion of a high number of ambiguous base pairs (Additional file 1: Supplementary Methods).

### Comparing binning performance and statistics

We applied computational (MetaBat) (21) and conformational capture methods (ProxiMeta Hi-C) (22) in order to bin assembled contigs into clusters that closely resembled the actual genomic content of unique species of rumen microbes (Additional file 1: Supplementary Methods). The number of contigs per bin varied based on the binning method; however, the long-read assembly bins had nearly an order of magnitude fewer contigs per bin than the short-read assembly regardless of method (Fig 2a). We also saw a clear discrepancy between binning methods, with ProxiMeta preferably binning smaller (< 2,500 bp) contigs with higher GC (> 42%) than MetaBat (Chi-Squared test of independence p < 0.001; Additional file: Fig S2).

**Figure 2.**
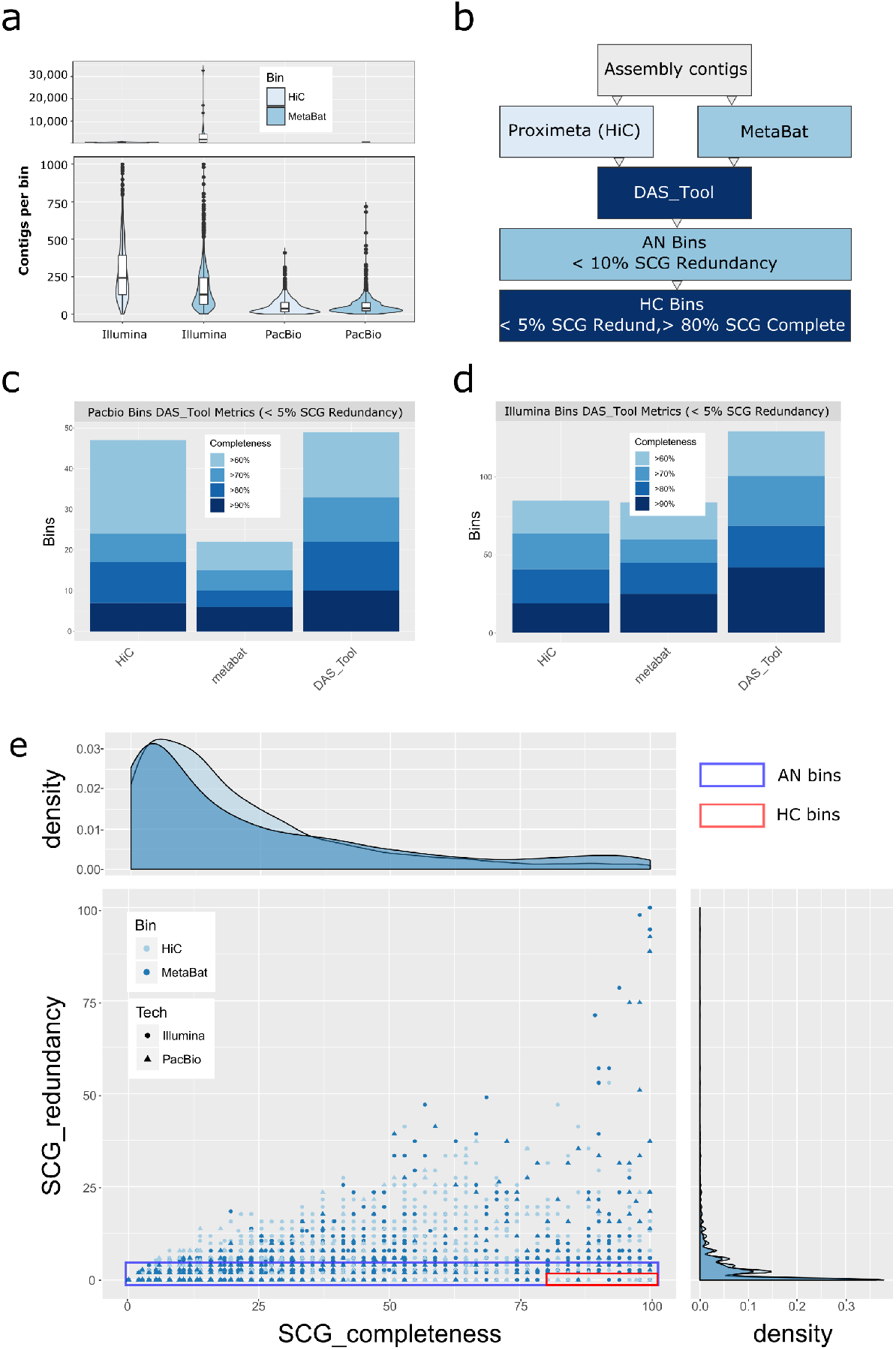
Identification of high quality bins in comparative assemblies highlights need for dereplication of different binning methods. (A). Binning performed by Metabat (light blue) and Proximeta Hi-C binning (Hi-C; blue) revealed that the long-read assembly consistently had fewer, longer contigs per bin than a short-read assembly. (B) Bin set division into Analysis (AN) and High Quality (HC) bins was based on DAS_Tool single copy gene (SCG) redundancy and completeness. Assessment of SCG completeness and redundancy revealed 22 and 48 high quality bins in the long-read (C) and short-read (D) assemblies, respectively. The Proximeta Hi-C binning method performed better in terms of SCG metrics in the long-read assembly. (E) Plots of all of identified bins in the long-read (triangle) and short-read (circle) assemblies revealed a wide range of chimeric bins containing high SCG redundancy. Bins highlighted in the blue rectangle correspond to the AN bins identified by the DAS_tool algorithm while the red rectangle corresponds to the HC bin set.

We further assessed bin quality and removed redundant contig-bin assignments between methods, using the single-copy gene (SCG) metrics of cluster contamination and completeness from the DAS_Tool (23) package (Fig 2c, d; Additional file 2, 3). We then sorted the revised DAS_Tool bins into a set of higher completion (HC) bins with less than 5% SCG redundancy and greater than 80% SCG completion, and a set of bins for analysis (AN) with less than 10% SCG redundancy (Fig 2b; Table 2). Since DAS_Tool assesses bin quality using bacterial and archaeal SCG metrics, we did not use a completion filter for the AN bin set as that would remove candidate high quality eukaryotic and viral bins from our analysis dataset. Our HC bin dataset contains 22 and 69 draft microbial genomes in the long-read and short-read datasets, respectively, with at least an 80% SCG completeness estimate and with less than 5% SCG redundancy (Fig 2e). The AN binset contained 1,028 and 3,757 long-read and short-read consolidated bins, respectively, which were used in subsequent analysis and characterization.

**Table 2.** Assembly bin taxonomic assignment and gene content

| Assembly | Bin Set | Avg # complete ORFs per contig<sup>2</sup> | Assembled Sequence Taxonomic Affiliation (Kbp)<sup>1</sup> |  |  |  |  |
| --- | --- | --- | --- | --- | --- | --- | --- |
|  |  |  | Archaea | Bacteria | Eukaryota | Viruses | No-Hits |
| Illumina | Unbinned | 1.10 | 25,843 | 1,614,799 | 50,562 | 4,280 | 676,394 |
|  | AN | 1.82 | 26,161 | 2,273,837 | 79,804 | 2,083 | 357,193 |
|  | HC | 6.53 | 1,101 | 116,800 | 910 | 4 | 1,172 |
| PacBio | Unbinned | 15.43 | 13,382 | 1,024,348 | 9,031 | 2,340 | 27,323 |
|  | AN | 15.79 | 7,096 | 771,677 | 6,083 | 1,016 | 12,289 |
|  | HC | 36.08 | 1,827 | 45,204 | 569 | 0 | 51 |

<sup>1</sup> Superkingdom taxonomic affiliation was based on contig-level assignments derived from the BlobTools/DIAMOND workflow.
<sup>2</sup> Complete ORFs were defined as Prodigal predictions that had a “partial” status of “00”, which indicates the presence of a start and stop codon for the ORF.

### Taxonomic classification reveals assembly bias

Taxonomic classification of the HC bin and AN binsets revealed a heavy preference towards the assembly of contigs of bacterial-origin vs archaeal- and eukaryotic-origin (Fig 3c; Additional file 1: Figures S3, S4). Both the short- and long-read HC bins contain only one bin, each consisting of Archaeal-origin sequence. The short-read archaeal HC bin was best classified as being a high quality draft from the *Thermoplasmatales* order; however, the long-read archaeal bin was identified as belonging to the genus *Methanobrevibacter* from the family *Methanobacteriaceae*. Contig taxonomic assignment generated by the BlobTools(24) workflow varied greatly among the short-read HC bins, with an average of 5 different phyla assignments per contig per bin (Additional files 4, 5). We identified 30 full-length (> 1,500 bp) predicted 16S rDNA genes in the HC bins, and only fragmentary (< 1,500 bp) 16S genes in the short-read assembly (Additional file 6). The long-read AN bins contained 239 full-length 16S genes, and all but 5 of the genes matched the original superkingdom taxonomic classification of the bin that contained the gene. Of these five discrepancies, three contigs were classified as “Eukaryotic” in origin, yet contained a predicted Archaeal 16S gene.

**Figure 3.**
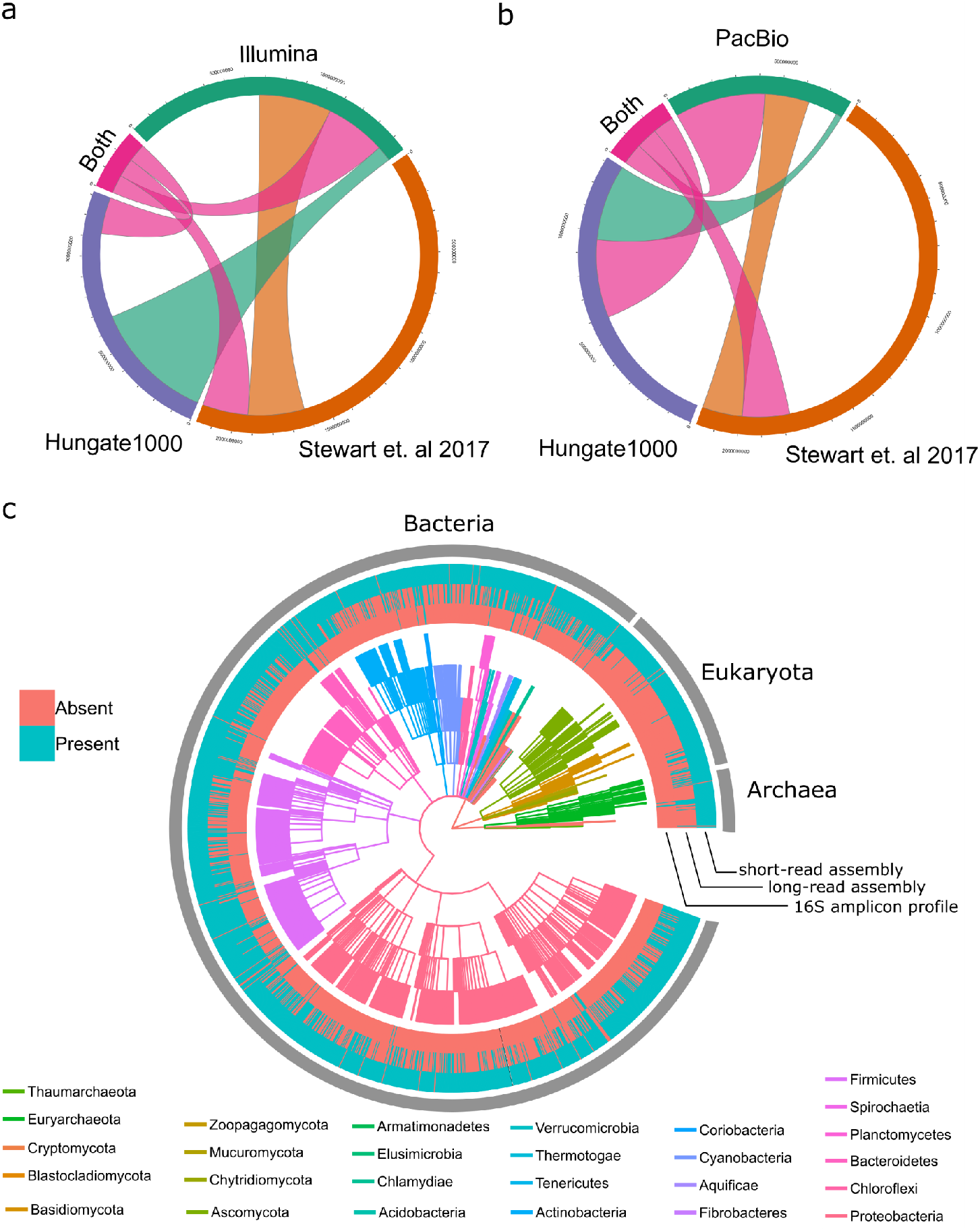
Dataset novelty compared to other rumen metagenome assemblies. Chord diagrams showing the contig alignment overlap (by base-pair) of the short-read (A) and long-read (B) AN bins to the Hungate1000 and Stewart et al. 2017 rumen microbial assemblies. The “Both” category consists of alignments of the short-read and long-read AN bins that have alignments to both Stewart et al. 2017 and the Hungate1000 datasets. (C) A dendrogram comparison of dataset sampling completeness compared to 16S V4 amplicon sequence data analysis. The outer rings of the dendrogram indicate presence (blue) or absence (red) of the particular phylotype in each dataset. Datasets are represented in the following order (from outer edge to internal edge): (1) the short-read assembly contigs, (2) the long-read assembly contigs, (3) and 16S V4 amplicon sequence data. The internal dendrogram represents each phylum in a different color (see legend), with individual tiers corresponding to the different levels of taxonomic affiliation. The outermost edge of the dendrogram consists of the genus-level affiliation.

### Comparison to other datasets reveals novel sequence

Contig novelty was assessed via direct overlap with other rumen metagenomic assemblies and via alignment with WGS reads from other publically accessible sources (Fig 3a,3b). We identified many contigs in our short-read and long-read assemblies that did not have analogous alignments to the recently published Stewart et al. (17) and Hungate 1000(18) assemblies. From our HC bins, 3697 and 92 contigs from the short- and long-read assemblies, respectively, did not align to any sequence in these two datasets, consisting of 23.5 and 1.7 megabases of assembled sequence that was missing from the previous, high quality, reference datasets for the rumen microbiome (Additional files 7, 8). Expanding the comparison to the AN binset, we identified 207,599 (669 Mbp) and 12,421 contigs (137 Mbp) in the short- and long-read assemblies, respectively, that did not have analogs in the previous rumen datasets (Fig 3a, 3b). From the AN bins with no alignments to other published datasets, we identified 152,739 and 185 contigs in the short- and long-read AN binsets that did not have analogous alignments to the other respective dataset (e.g. short-read vs long-read). This represented 435 Mbp of exclusive sequence in the short-read dataset not contained in our long-read dataset. However, we also identified 1.18 Mbp that was novel to the long-read AN bins despite the coverage disparity between the two datasets. Contigs that were exclusive to the long-read dataset were primarily of Firmicutes-origin, and had a higher median GC% value than other contigs in the long-read dataset (Kolmogorov-Smirnov p = 4.12 x 10^−5^).

We wanted to compare the short-read sequence of our sample against other published rumen WGS datasets to see if there were differences in sample community composition that may have accounted for novel assembled sequence in our dataset. We aligned the WGS reads from each dataset to our assemblies (Additional file 1: Table S15), and we created a normalized read depth matrix from contigs from our short- and long-read datasets that had at least a Genus-level taxonomic assignment. We then counted the number of times per Genus where a contig had a higher mapping percentage in our sample compared to all other sampled WGS datasets and used a hypergeometric test to calculate the relative enrichment of observations per taxonomic group. In both the short-read and long-read datasets, six AN bins belonging to the Eukaryote superkingdom were significantly enriched (hypergeometric p value < 1 x 10^−7^ in all cases), suggesting that the WGS reads derived from the SRA datasets had lower coverage of these fungal and protist genomes than our WGS reads. In terms of assembly-specific enrichment, the short-read assembly had a larger proportion of Eukaryote-origin contigs that were found to be significantly enriched in coverage compared to the long-read assembly (Additional file 9). This may have resulted from the previously noted assembly discrepancy, where the short-read assembly was far more likely than the long-read assembly to assemble low GC% Eukaryote contigs from lower coverage data despite sampling proportionally fewer reads from lower GC% tranches.

### Increased long-read contiguity results in more predicted ORFs per contig

We sought to assess whether the increased contiguity of the long-read assembly contigs provided tangible benefits in the annotation and classification of open reading frames (ORFs) in our AN bin dataset. From prodigal (25) annotation of the AN bins from both assemblies, we identified 868,429 and 984,623 complete ORFs in the long-read and short-read assemblies, respectively (Additional files 10, 11). We found a lower fraction of identified partial ORFs in the long-read AN bins (55,002 partial ORFs; 6% of the complete ORF count) compared to the short-read AN bins (365,281 partial; 37% of the complete ORF count). This would suggest that, despite a lower total count of total ORFs identified, the long-read bins more frequently contained complete ORFs than did the short-read bins. We also found a higher mean count of ORFs per contig in the long-read AN bins (mean: 15.79) than the short-read bins (mean: 2.69). This difference in average counts was found to be significant (Kolmogorov-Smirnov test p value < 0.001) and may be due to the presence of longer contigs found in the long-read assembly dataset. The majority of partial ORF predictions occur within the first 50 bp of contigs in the long read (99.9%; Chi squared p < 0.001) and short read (95.2%; p < 0.001) AN bins, suggesting that ORFs were prematurely terminated by contig breaks. In the short read AN bins, a surprising proportion of ORFs missing both a start and stop codon (23,458 ORFs; 6.4% of the total count of partial ORFs) occur near the beginning of the contig compared to the long read bin set (56 ORFs). However, we identified a slight discrepancy in ORF length between the long-read (median ORF length: 533 bp) and short-read (median: 584 bp) assemblies, with the later containing longer predicted ORFs than the long-read assembly.

We identified clear differences in gene content between the two assemblies that suggest a bias in functional ORF classification and discovery. Using cluster of orthologous group (COG) assignments to predicted ORFs in both assemblies, we identified a discrepancy in the proportional count of several major COG categories. The long-read Bacterial AN binset contained proportionally more L (Replication, recombination and repair), Q (Secondary metabolites), P (Inorganic ion transport) and H (Coenzyme transport/metabolism) COG ORFs than the short-read bins, and proportionally less J (Translation) and M (Cell wall) COG ORFs (Additional file 1: Figure S5, Table S11) as determined by a Fisher’s exact test. Conversely, the long-read Archaeal AN bins contained more J and C (Energy production and conversion) COG ORFs than the short-read assembly, while still having proportionally fewer M COG ORFs. The V (Defense mechanisms) COG category was proportionally higher in the short-read assembly for both the Bacterial and Archaeal lineages, suggesting a higher proportion of defense-related ORFs were assembled in that dataset.

### Host-prophage association and CRISPR array identification

Longer reads have the potential to provide direct sequence-level confirmation of prophage insertion into assembled genomes by spanning direct repeats that typically flank insertion sites (26). To identify candidate host-specificity for assembled prophage genomes, we used a heuristic alignment strategy with our error corrected long-reads (Additional file 1: Supplementary Methods) and Hi-C intercontig link density calculations. PacBio sequence data have a known propensity for chimerism(27); however, we assumed that identical, chimeric PacBio reads would be unlikely to be seen more than once in our dataset. Similarly, we filtered Hi-C read alignments to identify virus-host contig pairs with higher link counts to identify host-viral associations in each assembly (Additional file 1: Supplementary Methods). Several viral contigs in the long-read assembly had substantial associations with contig groups affiliated with more than one genus (a maximum of 11 distinct genus-level classifications for one viral contig from the Myoviridae), suggesting a wide host-specificity for these species (Fig 4a). Long-read assembly viral contigs with multiple candidate host associations were identified as belonging to the Podoviridae, Myoviridae and Siphoviridae families, which are viral families typically encountered in bovine rumen microbial samples (28). Viral contigs from the short-read assembly were associated with fewer candidate host genus OTUs (four distinct associations at maximum; Fig 4b). It is possible that the shorter length of Illumina assembly viral contigs (average size: 4140 bp, standard deviation(sd): 5376 bp) compared with the long-read assembly contigs (average: 20,178bp, sd: 19,334 bp) may have reduced the ability to identify host-phage associations in this case. Having identified read alignments between viral contigs and non-viral contigs, we sought to leverage conformational capture via Hi-C to see if we could confirm the viral-host associations.

**Figure 4.**
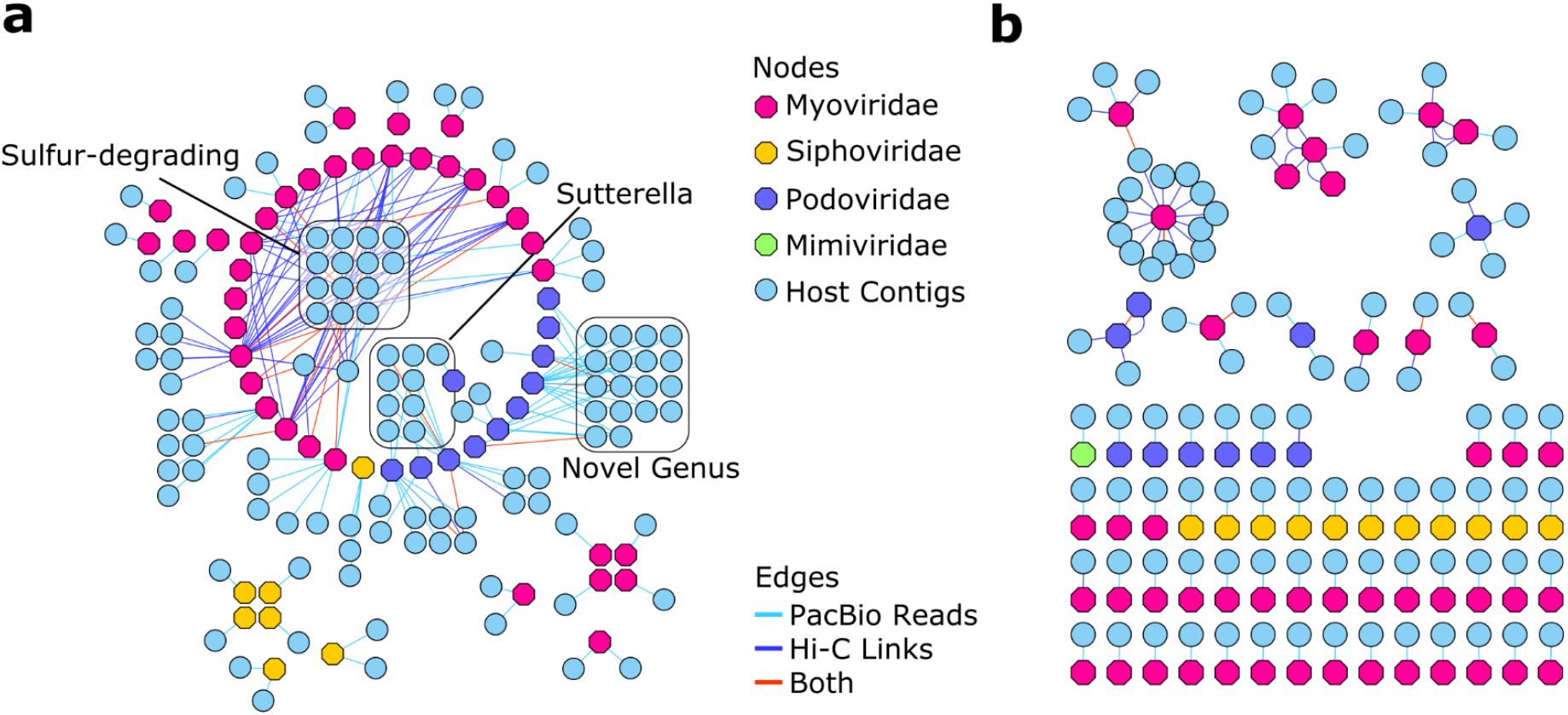
Network analysis of long-read alignments and Hi-C inter-contig links identifies hosts for assembled viral contigs. In order to identify putative hosts for viral contigs, PacBio read alignments (light blue edges) and Hi-C inter-contig link alignments (dark blue edges) were counted between viral contigs (hexagons) and non-viral contigs (circles) in the long-read assembly (A) and the short-read assembly (B). Instances where both PacBio reads and Hi-C inter-contig links supported a viral-host assignment are also labeled (red edges). The long-read assembly enabled the detection of more viral host-associations in addition to several cases where viral contigs may display cross-species infectivity. We identified several viral contigs that infect important species in the rumen, including those from the genus *Sutterella*, and several species that metabolize sulfur. In addition, we identified a candidate viral-association with a novel genus of rumen microbes identified in this study.

We found that our Hi-C link analysis and PacBio read alignment analysis had very little overlap; however, we identified a tendency for each method to favor a different class of virus-host association which suggested that the methods were complementary rather than antagonistic (Additional file 12). Approximately 10% (long-read: 19 out of 188 pairs; short-read: 6 out of 109) of the host-viral contig associations had supporting evidence from both PacBio read alignments and Hi-C intercontig links. In nearly all highly-connected viral contig pairs (greater than two additional contig associations) we observed evidence of host specificity from both methods even if it was for different host contigs. We also identified a bias in the host-viral family associations, where putative hosts for the Myoviridae were more likely to be identified via Hi-C than other viral families (Fig 4a). Myoviridae family viral specificity for the sulfur-reducing *Desulfovibrio* and the sulfur-oxidizing *Sulfurovum* genera were primarily identified through Hi-C contig links (Fig 4a, box: “Sulfur-degrading”). However, viral associations between the *Sutterella* and a previously unreported genera of rumen bacteria were primarily identified via PacBio read alignments and had little Hi-C intercontig link support.

We also tested the ability of longer read sequence data to resolve highly repetitive bacterial defense system target motif arrays, such as those produced by the CRISPR-Cas system, in our dataset. Despite having less than one third of the coverage of the short-read dataset, our long-read assembly contained two of the three large CRISPR arrays (consisting of 105 and 115 spacers, respectively) in our combined assembly dataset (Fig 5a). The short-read dataset (597 CRISPR arrays) contained approximately five-fold more identifiable CRISPR arrays than the long-read dataset (122 arrays), which is commensurate with the difference in the size of each assembly (5 Gbp vs 1 Gbp, respectively).

**Figure 5.**
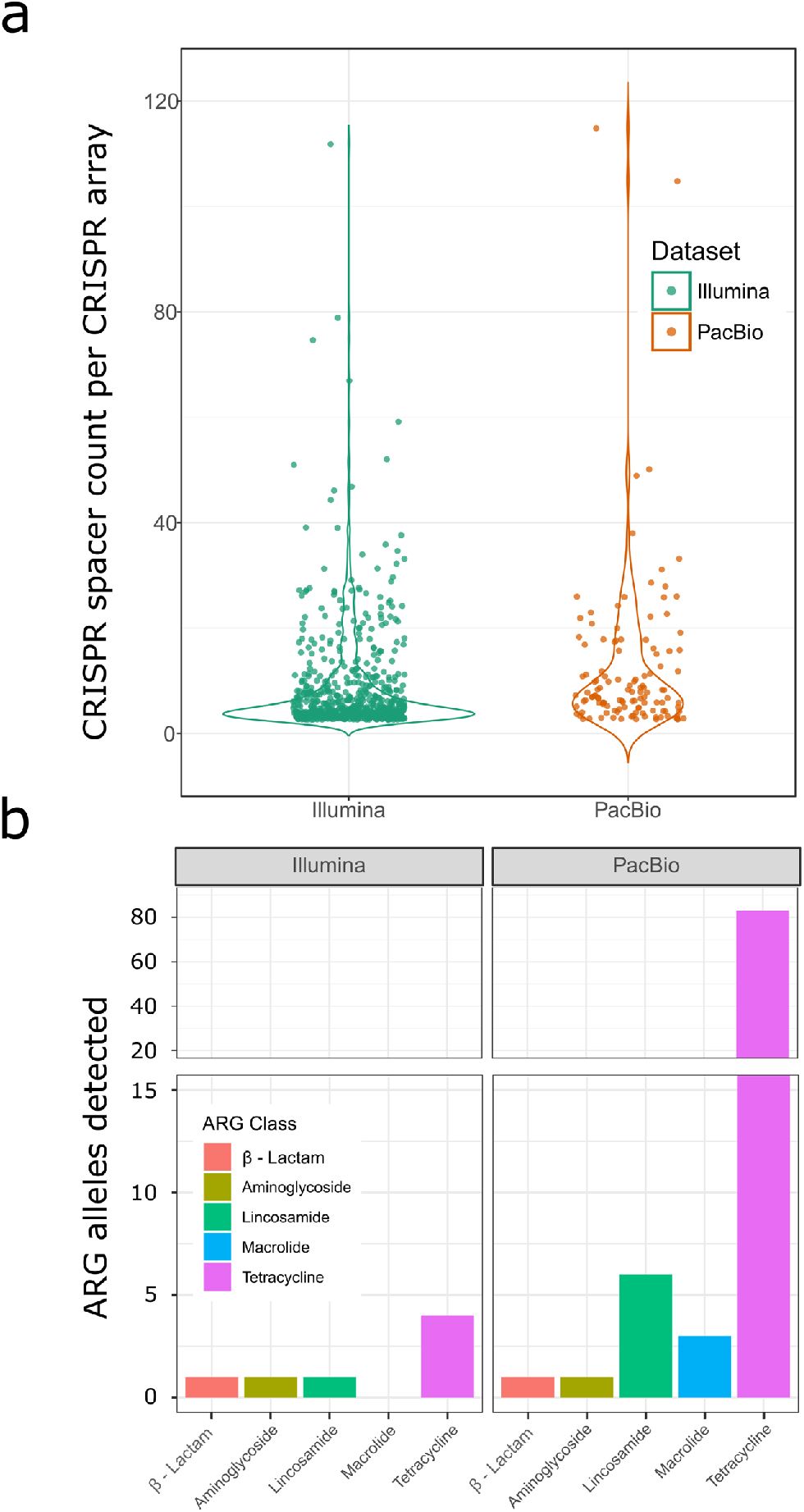
CRISPR array identification and ARG allele class counts were influenced by assembly quality. (A) The long-read assembly (dark orange) contigs had fewer identified CRISPR arrays than the short-read contigs (dark green); however, the CRISPR arrays with the largest count of spacers were overrepresented in the long-read assembly. (B) The long-read assembly had 13-fold higher anti-microbial resistance gene (ARG) alleles than the short-read assembly despite having 5-fold less sequence data coverage. The Macrolide, Lincosamide and Tetracycline ARG classes were particularly enriched in the long-read assembly compared to alleles identified in the short-read assembly.

### Antimicrobial resistance gene detection

Due to the frequent use of antibiotics in livestock production systems to treat disease and improve production, we wanted to assess the utility of longer-reads in detecting novel ARG alleles in assembled microbial genomes (Fig 5b). The long-read assembly (ARG allele count: 94) was found to contain over an order of magnitude more identifiable ARG alleles than the short-read assembly (ARG allele count: 7), despite the major coverage discrepancies between the two datasets. The major contributor to this discrepancy was found in the Tetracycline resistance gene class, as the long-read assembly contained 80 ribosomal protection and 3 efflux ARGs that are predicted to confer tetracycline resistance. By contrast, only 2 ribosomal and 2 efflux Tetracyline ARGs were identified in the short-read assembly. Using the contigs containing these ARG alleles as anchors in our alignment of Hi-C read pairs, we attempted to identify horizontal transfer of these alleles using Hi-C intercontig link signal (Additional file 1: Supplementary Methods). We identified clusters of *Prevotella* bins, and clusters of bins from the Clostridiales and Bacteroidales that higher contig-link density with ARG allele contigs in our dataset (Additional file 1: Figure S6; Additional file 13). These associations may represent potential horizontal transfer of these alleles; however, we note that inter-contig link density was relatively low in our comparisons (average alignments density was less than 2 reads per pair) and that ambiguous alignment to orthologous sequence could present false-positive signal in this analysis.

## Discussion

Whole metagenome shotgun sequencing and assembly has often relied exclusively on short-read technologies due to the cost-effectiveness of the methods and the higher throughput that they provide. While such strategies are often able to efficiently generate sufficient read depth coverage to assemble fragments of organisms in the community, we demonstrate that biases inherent in singular technologies suitable for metagenome assembly result in an incomplete assembly/binning of the actual community. For example, we exclusively assembled a member of the Archaeal order *Thermoplasmatales* in our short-read dataset and a member of the Archaeal genus *Methanobrevibacter* in the long-read assembly. Several taxonomic studies using short-read 16S-based methods have shown that the CO_2_-reducing *Methanobrevibacter* are one of the most abundant genera of methanogenic Archaea in the rumen (29), which was not reflected in our short-read assembly dataset despite higher depths of coverage. Conversely, we found that the short-read assembly was better at resolving genomic fragments of the Eukaryotic Superkingdom, which were relatively underrepresented in the long-read assembly. Given that we sequenced the same biological sample in all of our analyses, these discrepancies suggest that each technology samples different portions of the rumen microbial community. Our data suggest that each technology’s unique purview can be attributed to compositional differences of the genomes among taxonomic superkingdoms (Fig 1c), genomic GC% (Fig 1b), and the presence of mobile DNA (Fig 4, Additional file 1: Figure S6).

We identified a GC% bias in our short-read data relative to our long-read reads; however, this relative bias was reversed in comparisons of the GC content of the final assemblies, where our short-read assembly had more -- albeit shorter -- assembled contigs in lower GC% tranches (Fig 1b). These differences are most likely due to the different error rates and degrees of coverage of reads from the two sequencing technologies and the algorithms used by the different assembly programs to correct for errors. Paradoxically, the short-read assembly sampled proportionally fewer reads at higher and lower GC tranches, but was able to incorporate even fragmentary information from these tranches into smaller contigs. The long-read assembly, by contrast, required sufficient coverage of reads to appropriately correct for errors and this meant that many lower GC% reads were discarded due to assembly constraints, as we demonstrate in our read alignment overlap analysis (Additional file 1: Figure S1). Protists may represent a large proportion of this lower GC% community, and their genomes likely consist of highly repetitive sequence that would require higher depths of long-read coverage to sufficiently traverse (20). The use of improved error-correction methods or circular-consensus sequence reads (30,31) are likely to provide substantial benefits for downstream annotation and may enable the assembly of the low-abundance, low-GC% species that were poorly represented in our long-read assembly. Regardless, we found that even a lower depth of coverage of high error-rate long-reads better resolved biological features in the highly abundant strains than those detected in our short-read assembly.

We identified many biological features in our sample that would be missed if only a single technology/method was used for each step of the assembly, binning and analysis of our dataset. While differences in DNA sequencing technology represented a far smaller proportion of the total missing sequence (~ 0.5%) in our pairwise comparison, the missing fraction consisted of relatively large contigs with SCGs suitable for binning, as well as 7,886 complete ORFs. Larger contigs in the long-read dataset also resulted in a higher average count of annotated ORFs per contig than the short-read dataset by a factor of seven. This contiguity of gene regions is particularly important in bacterial classification, where functional genes of particular classes can be arranged in complete and phased operons. It is highly likely that this increase in contiguity contributed to the massive discrepancy in ARG allele identification between the two assemblies. We noted a significant increase in detected Tetracycline resistance alleles in our long-read assembly of a rumen metagenome from a concentrate-fed animal, which contradicts previous work using short-read assemblies that found that animals fed concentrates should have few Tetracycline resistance alleles (32). Calves in the sampled research herd (UW-Madison, Dairy Forage Research Center) are given Chlortetracycline during inclement weather and Tetracycline is applied topically to heel warts on adult animals. It is possible that incidental/early exposure to this antibiotic has enabled the proliferation of tetracycline resistance alleles in the rumen community, and this proliferation was only detected in our long-read assembly. Previous studies have demonstrated the benefit of using longer reads in ARG allele –associated satellite DNA tracking (33) and ARG allele amplicon sequencing (34). To our knowledge, this is the first survey to identify the benefits of long-reads in de novo assembly of ARG alleles from a complex metagenomics sample.

We also identified discrepancies between our selected computational (MetaBat) and proximity ligation (ProxiMeta Hi-C) binning methods that suggest that a combination of binning techniques are needed to identify all complete MAGs in a metagenomic sample. Contig binning comparisons suggest that MetaBat successfully binned contigs from the low-GC% contig tranches; however, it failed to incorporate the same proportion of smaller contigs in bins from the short-read (< 2,500 bp) or long-read (< 10,000 bp) assemblies as the ProxiMeta method. Smaller contigs most likely result from low-sequencing coverage regions or high copy orthologous genomic segments in a metagenomic sample. Both of these problems may have confounded the tetranucleotide frequency and coverage depth estimates used by MetaBat to bin our contigs, resulting in their lower frequencies in that binset. We did note some issues in DAS_tool dereplication of our dataset, where DAS_tool may have aggressively pruned contigs from MetaBat bins. However, our data suggests that MetaBat may have included far more contamination due to cross-Kingdom SCGs, thereby resulting in this aggressive filtration.

In order to identify the horizontal transfer of mobile DNA in the rumen, we exploited two technologies to identify candidate hosts for transferred ARG alleles and assembled viral contigs. We observed inter-contig link associations between ARG allele contigs and bins that consisted of species from the Clostridiales and Bacteroidales. Evidence of identical ARG allele orthologs belonging to both classes was previously found in human colon samples (35); however, we note that our analysis shows only a precursory association of the context of identified ARG alleles and prospective host bins. We were unable to identify the exact vector that may enable the cross-species transfer of several of these alleles, but we suspect that lateral transfer of ARG alleles may be an adaptation of rumen bacterial species against antibiotic challenge as noted above. Direct evidence of the horizontal transfer of mobile elements was observed in identified novel host-viral associations that we detected by using a combination of PacBio long-read alignments and Hi-C intercontig link analysis. Proximity ligation has been previously used to detect host-virus associations (36); however, our combination of technologies potentially reveals new insights in the biology of the interaction between host and phage. We found a clear preference between the two methods in the detection of viral family classes, with Hi-C intercontig links preferring the Myoviridae viral family and our PacBio read alignments preferring all other viral families. This preference may reflect the nature of the activity of these viruses, as some genera of the Myoviridae family are known to have short lytic cycles (37) as opposed to long-term lysogenic life-cycles found in other viral families. We also identified viral-host association with several contigs within bins identified as belonging to the *Desulfovibrio* and *Sulfurovum* genera. Viral auxiliary metabolic genes related to sulfur metabolism were previously identified in assembly of rumen viral populations (28), and our study may provide a link to the putative origins of these auxiliary genes in host genomes that are known to metabolise sulfur compounds. We identified two ORFs annotated as 3′-Phosphoadenosine-5′-phosphosulfate (PAPS) genes in a viral contig in the long-read assembly that was associated with host contigs assigned to the *Dehalococcoides*. We did not detect any auxiliary metabolic genes in the short-read assembly. Additionally, the short-read assembly served as the basis of fewer host-viral contig associations in both Hi-C and PacBio read analyses, suggesting that assembled short-read viral contigs may have been too small or redundant to provide a useful foundation for alignment-based associations.

We recommend that future surveys of complex metagenomic communities include a combination of different DNA sequencing technologies and conformational capture techniques (ie. Hi-C) in order to best resolve the unique biological features of the community. If our analysis was restricted to the use of the short-read WGS data and one computational binning technique (MetaBat), we would have missed 139 out of 250 of the top dereplicated DAS_Tool short-read bins contributed by ProxiMeta binning. Our long-read dataset further contributed 7,886 complete ORFS, 97 ARG alleles and 188 host-virus associations, with Hi-C signal providing further evidence of host-virus associations. We demonstrate that even a small proportion of long-reads can contribute high quality metagenome bins, and that the long-read data provided by the technology is suitable for uncovering candidate mobile DNA in the sample. We also note that the inclusion of a computational binning method (Metabat) with a physical binning technique (ProxiMeta; Hi-C) further increased our count of high quality, DAS_Tool dereplicated bins, likely due to each method sampling a different pool of organisms. Therefore, the DAS_Tool dereplication of both sets of bins increased our final counts of high quality (> 80% completion) bins by 30-60% in the long-read and short-read assemblies. If a metagenomics WGS survey is cost-constrained, our data suggests that a computational method, such as MetaBat, currently cannot fully compensate for the GC% bias and repetitive, orthologous DNA issues that could reduce the completeness of a downstream short-read assembly. Still, we suspect that such projects will be able to assemble and characterize the abundant, moderate-GC portion of the metagenome community sufficiently for analysis.

Further refinements could improve characterization of the rumen microbial community and other complex metagenomic communities in general. For example, microbes present in low abundance (or transient species) still represent a challenge to all of the technologies used in our survey. A sample fractionation method similar to one used by Solden et al. (38) would enable better, targeted coverage of these communities in future surveys while losing the ability to determine relative abundance estimates for strains. In the absence of targeted sample enrichment, co-assembly with other sampled datasets (17), low-error rate long-reads (31) or real-time, selective read sequencing (39) would enable sampling of lower abundant strains. Additionally, there is a need for a rigorous method to combine and/or scaffold metagenome assemblies with high-error long-reads. Our attempts to combine our short-read and long-read datasets using existing scaffolding and assembly software failed to produce a significant improvement in assembly contiguity and quality. The complexity of the data will likely require a specialized solution that can also resolve issues that result from excessive strain heterogeneity.

## Conclusions

We demonstrate the benefits of using multiple sequencing technologies and proximity ligation in identifying unique biological facets of the cattle rumen metagenome and we present data that suggests that each has a unique niche in downstream analysis. Our comparison identified biases in the sampling of different portions of the community by each sequencing technology (e.g. bias in GC% representation), suggesting that a singular DNA sequencing technology is insufficient to characterize complex metagenomic samples. Using a combination of long-read alignments and proximity ligation, we identified putative hosts for assembled bacteriophage at a resolution previously unreported in other rumen surveys. These host-phage assignments support previous work that revealed increased viral predation of sulfur-metabolising bacterial species; however, we were able to provide a higher resolution of this association, identify potential auxiliary metabolic genes related to sulfur metabolism, and identify phage that may target a diverse range of different bacterial species. Furthermore, we found evidence to support that these viruses have a lytic lifecycle due to a higher proportion of Hi-C inter-contig link association data in our analysis. Finally, it appears that there may be a high degree of mobile DNA that was heretofore uncharacterized in the rumen, and that this mobile DNA may be shuttling antimicrobial resistance gene alleles among distantly related species. These unique characteristics of the rumen microbial community would be difficult to detect without the use of several different methods and techniques that we have refined in this study, and we recommend that future surveys incorporate these techniques to further characterize complex metagenomic communities.

## Methods

### Sample selection, DNA extraction and Hi-C library preparation

Rumen contents from one multiparous Holstein cow housed at the University of Wisconsin, Madison, campus were sampled via rumen cannula as previously described (40). The sampled cow was in a later period of lactation and was being fed a total mixed ration. Rumen solids and liquids were combined in a 1:1 volume mix, and then were agitated using a blender with carbon dioxide gas infusion as previously described (40). DNA was extracted via the protocols of Yu and Morrison (41) albeit with several modifications to the protocol to increase yield. To improve DNA precipitation, an increased volume of 10 M ammonium acetate (20% of the supernatant volume) was added. Additionally, DNA pellets were not vacuum dried so as to reduce the potential for single-strand nicking due to dehydration. DNA quality was assessed via Fragment Analyzer spectra and spectrophotometric assays.

Different DNA extraction methods can result in substantial observed differences in strain- and species-level assignments depending on the recalcitrance of the cell wall of individual cells (8). However, contemporary long-read sequencing platforms require input DNA to be devoid of single-strand nicks in order to maximize sequence read lengths (42). Indeed, our observed, average subread length for the long-read dataset was almost half (7,957 ± 4,957 bp) the size of our original Fragment Analyzer spectra peaks (~ 14,651 bp), suggesting that the bacterial cell lysis still impacted DNA molecule integrity (Additional file 1: Figure S8). Regardless, the average subread length was 9 kb and we were able to sequence a total of 52.92 gigabases of raw PacBio data for our downstream analysis.

Portions of the rumen contents samples were fixed by a low concentration formaldehyde solution before DNA extraction as previously described (43). Fixed samples were subject to the same DNA extraction protocol as listed above, processed by Phase Genomics (Seattle, WA) and sequenced on a HiSeq 2000.

### Long-read and short-read DNA sequencing

Tru-seq libraries were created from whole DNA preps for the sample as previously described (44). Samples were run on a single Illumina NextSeq500 flowcell using a 300 cycle SBS kit to produce 150 bp by 150 bp paired-end reads.

DNA samples from each cow were size selected to a 6 kb fragment length cutoff using a Blue Pippen (Sage Science; Beverly, MA). Libraries for SMRT sequencing were created as previously described (6) from the size-selected DNA samples. We generated 7.57 and 45.35 Gbp of PacBio uncorrected reads using the PacBio RSII (8 cells) and PacBio Sequel (21 cells), respectively. A total of 52.92 Gbp of subread bases with an average read length of 6623.33 bp were generated on all samples using PacBio sequencers (Additional file 1: Table S14).

### Genome assembly and binning

PacBio raw reads were assembled by Canu v1.6+101 changes (r8513). We ran five rounds of correction to try to recover lower-coverage reads for assembly using the parameters “correct corMinCoverage=0 genomeSize=5m corOutCoverage=all corMhapSensitivity=high”. The input for each subsequent round were the corrected reads from the previous step. Finally, the assembly was generated via the parameters “-trim-assemble genomeSize=5m oeaMemory=32 redMemory=32 correctedErrorRate=0.035”. The assembly was successively polished twice with Illumina data using Pilon restricted to fix indel errors using the “-fix indels” and “-nostrays” parameters. Pilon correction was automated using the slurmPilonCorrectionPipeline.py script available at the following repository: https://github.com/njdbickhart/RumenLongReadASM. We generated a second set of PacBio corrected reads for the viral-association and GC-read overlap analyses using the options “-correct corMinCoverage=0 genomeSize=5m corOutCoverage=all corMhapSensitivity=high corMaxEvidenceCoverageLocal=10 corMaxEvidenceCoverageGlobal=10” to restrict the global filter to avoid over-smashing similar sequences during correction. Illumina reads were assembled using MegaHit v1.1.2 using parameters --continue --kmin-1pass -m 15e+10 --presets meta-large --min-contig-len 1000 -t 16 and otherwise default settings.

Reads from other rumen WGS datasets (Additional file 1: Table S15) were aligned to assembled contigs from both assemblies with BWA MEM(45) and were used in Metabat2 binning(21). Metabat2 was run with default settings using the coverage estimates from all rumen WGS datasets (Additional file 1: Supplementary methods). Hi-C reads were aligned to assembled contigs from both assemblies using BWA MEM (45) with options -5S, and contigs were clustered using these alignments in the Phase Genomics ProxiMeta analysis suite(43). We noted a difference in bin contamination between the two methods, where Metabat tended to have more bins with greater than 10% CheckM(46) Contamination (76 out of 1347 short-read bins) compared to the ProxiMeta bins (29 out of 3,664 bins; Chi-Squared p < 0.001).

Using the ProxiMeta and MetaBat bin assignments as a seed, we consolidated assembly bins for each assembly using the DAS_Tool pipeline (23). The dereplication algorithm of DAS_Tool modifies input bin composition in an iterative, but deterministic, fashion, so we also validated the quality of our input bins by using CheckM (46) quality metrics in addition to the DAS_Tool SCG metrics (Fig 2c, 2d). We noted some discrepancies in the CheckM quality metrics and those estimated by DAS_Tool for our input and dereplicated MetaBat bins, respectively (Additional file 1: Figure S9, S10). CheckM tended to overestimate the quality of MetaBat input bins and dereplicated bins in each assembly, which may have due to the inclusion of proportionally more cross-Kingdom SCGs in the MetaBat bins as assessed by DAS_Tool. As a result, DAS_Tool dereplication was far more permissive at removing bins from our MetaBat dataset (average 69 +/− 204 contigs removed per bin) than our ProxiMeta dataset (average 23 +/− 30 contigs) in our short-read dataset. For further details on assembly binning and bin dereplication, please see Additional file 1: Supplementary Methods.

### Assembly statistics and contaminant identification

General contig classification and dataset statistics were assessed using the Blobtools pipeline (24). To generate read coverage data for contig classification, paired-end short read datasets from 16 SRA datasets and the Illumina sequence data from this study were aligned to each contig and used in subsequent binning and contaminant identification screens. For a full list of datasets and accessions used in the cross-genome comparison alignments, please see Additional file 1: Table S15. Assembly coverage and contig classifications were visually inspected using Blobtools (24). Comparisons between assembled contigs and other cattle-associated WGS metagenomics datasets were performed by using MASH (47) sketch profile operations and minimap2(48) alignments. Datasets were sketched in MASH by using a kmer size (-k) of 21 with a sketch size of 10,000 (-s). Minmap2 alignments were performed using the “asm5” preset configuration. DIAMOND (49) alignment using the Uniprot reference proteomes database (release: 2017_07) was used to identify potential taxonomic affiliation of contigs through the Blobtools metagenome analysis workflow (24). MAGpy (50) was also used to suggest putative names for the short and long read bins. CheckM (46) version 1.0.11 was used to assess bin contamination and completeness separately from the DAS_Tool SCG quality metrics.

### ORF prediction, gene annotation and taxonomic affiliation

Open reading frames were identified by Prodigal (25) (v 2.6.3) as part of the DAS_Tool pipeline. Gene ontology (GO) term assignment was performed using the Eggnog-mapper pipeline (51) using the same Diamond input alignments used in the Blobtools analysis.

Assembly bin functional classification was determined using the FAPROTAX workflow (52), using the Uniprot/Diamond/Blobtools-derived taxonomy of each contig. In order to deal with uncertain species-level classifications for previously unassembled strains, taxonomic affiliations were agglomerated at the genus level for dendrogram construction. The reference tree was created from NCBI Common Tree (https://www.ncbi.nlm.nih.gov/Taxonomy/CommonTree/wwwcmt.cgi) and plotted in the R package ggtree (53).

### Viral-host association prediction and Hi-C intercontig link analysis

In order to identify potential virus-host links, we used a direct long-read alignment strategy (PacBio alignment) and a Hi-C intercontig link analysis (Hi-C). Briefly, contigs identified as being primarily viral in-origin from the Blobtools workflow were isolated from the short-read and long-read assemblies. These contigs were then used as the references in an alignment of the error-corrected PacBio reads generated in our second round of Canu correction (please see the “Genome Assembly and Binning” section above). We used Minimap2 to align the PacBio dataset to the viral contigs from both datasets using the “map-pb” alignment preset. Resulting alignment files (“paf”) were subsequently filtered using the “selectLikelyViralOverhangs.pl” script, to selectively identify PacBio read alignments that extend beyond the contig’s borders. We then used the trimmed, unaligned portions of these reads in a second alignment to the entire assembly to identify putative host contigs (Additional file 1: Supplementary methods). A viral-host contig pair was only identified if two or more separate reads aligned to the same viral/non-viral contig pair in any orientation.

Hi-C intercontig link associations were identified from read alignments of the Hi-C data to each respective assembly. BAM files generated from BWA alignments of Hi-C reads to the assemblies were reduced to a bipartite, undirected graph of inter-contig alignment counts. The graph was filtered to identify only inter-contig links that involved viral contigs and that had greater than 20 or 10 observations in the long-read and short-read assembly, respectively. The information from both methods was combined in a qualitative fashion using custom scripts (Additional file 1: Supplementary methods). The resulting dataset was visualized using Cytoscape(54) with the default layout settings, or the “attribute circle” layout option depending on the degrees of viral-contig associations that needed to be visually represented.

### CRISPR-CAS spacer detection and ARG detection

ARG homologues were identified using BLASTN with the nucleotide sequences extracted from the Prodigal ORFs locations as a query against the transferrable ARG ResFinder database (55). Hits with a minimum 95% nucleotide sequence identity and 90% ARG sequence coverage were retained as candidate ARGs. Hi-C linker analysis identifying ARG gene contig-associations was derived from Proximeta bin data and Hi-C read alignments by counting the number of read pairs connecting contigs in each bin to each ARG. The procedure for identifying these associations was similar to the protocol used to identify Hi-C-based, Viral-Host associations. Briefly, a bipartite, undirected graph of inter-contig alignment counts was filtered to contain only associations originating from contigs that contained ARG alleles and had hits to non-ARG-containing contigs. This graph was then converted into a matrix of raw association counts, which were then analyzed using the R statistical language (version 3.4.4). Taxonomic affiliations of contigs were derived from Blobtools, whereas the taxonomic affiliations of AN bins were derived from ProxiMeta MASH (47) and CheckM(46) analysis.

### Ethics approval and consent to participate

All animal work was approved by the University of Wisconsin-Madison Institutional Animal Care and Use Committee under protocol A005590-A04. Research was conducted under an IACUC approved protocol in compliance with the Animal Welfare Act, PHS Policy, and other Federal statutes and regulations relating to animals and experiments involving animals. The facility where this research was conducted is accredited by the Association for Assessment and Accreditation of Laboratory Animal Care, International and adheres to principles stated in the Guide for the Care and Use of Laboratory Animals, National Research Council, 2011.

### Availability of data and materials

The datasets generated and/or analysed during the current study are available in the NCBI SRA repository under Bioproject: PRJNA507739. The assemblies, bins and other supplementary data are available at this URL: https://obj.umiacs.umd.edu/marbl_publications/rumen/index.html. A description of commands, scripts and other materials used to analyze the data in this project can be found in the following GitHub repository: https://github.com/njdbickhart/RumenLongReadASM

## Supporting information

## Acknowledgments

DMB was supported by USDA CRIS project 5090-31000-026-00-D. KB was supported by USDA NIFA AFRI grant 5090-31000-026-06-I, “Reassembly of Cattle Immune Gene Clusters for Quantitative Analysis.” KPB was supported by USDA CRIS project 5090-21000-064-00-D. TPLS was supported by USDA CRIS project 3040-31000-100-00-D. BJH and JVK were supported by USDA CRIS project 8042-32000-110-00-D. IL, STS, and MOP were supported by NIAID grant R44AI122654-02A1. SK and AMP were supported by the Intramural Research Program of the National Human Genome Research Institute, US National Institutes of Health. This work used the computational resources of the NIH HPC Biowulf cluster (https://hpc.nih.gov). GS was supported by a USDA NIFA AFRI Foundational grant 2015-67015-23246. The authors would like to thank Mark Boggess and Michael Maroney for helpful discussions.

## Author Contributions

DMB, MW, SK, AMP and TPLS conceived the project and designed the experiments. KPB and LMC collected the rumen sample and extracted the DNA. CH, TPLS, IL, MOP and STS created sequencing libraries and sequenced the sample. DMB, MW, SK and TPLS wrote the manuscript. All other authors contributed specific analysis that was included in the submitted manuscript.

## Competing Interests

CH is an employee of Pacific Biosciences. IL, MOP and STS are employees of Phase Genomics. The other authors declare no additional competing interests.

