## Supplementary material for "Assignment of virus and antimicrobial resistance genes to microbial hosts in a complex microbial community by combined long-read assembly and proximity ligation"

Table S1 (short-read assembly contigs and bin assignments): (separate file)

Table S2 (long-read assembly contigs and bin assignments): (separate file)

Table S3 (Illumina Taxonomic assignment tables): (separate file)

Table S4 (Pacbio taxonomic assignment tables): (separate file)

Table S5 (PacBio full-length 16S genes):

Table S6 (Illumina Novel sequence missing from previous rumen assemblies): (separate file)

Table S7 (Pacbio Novel sequence missing from previous rumen assemblies): (separate file)

Table S8 (Hypergeometric test for enrichment):

Table S9 (Illumina prodigal predictions): (separate file)

Table S10 (Pacbio prodigal predictions): (separate file)

Table S11. Enrichment comparison of COG/KOG annotated-genes between the short read and long read datasets for Archaea, Bacteria, and Eukaryota.

| **Ruminal Microbe** | | ***Archaea*** | | | ***Bacteria*** | | | ***Eukaryota*** | |
| --- | --- | --- | --- | --- | --- | --- | --- | --- | --- |
| **COG/KOG Category** | **Code** | ***Short Read*** | ***Long Read*** | ***Short Read*** | | ***Long Read*** | ***Short Read*** | | ***Long Read*** |
| **Information storage and processing** | | | | | | | | | |
| Translation, ribosomal structure and biogenesis | J | 3,512 (7.75%)† | 1,123 (10.13%) | 219,507 (7.1%)* | | 46,271 (6.02%) | 4,725 (6.72%)† | | 615 (10.32%) |
| RNA processing and modification | A | 6 (< 0.01%) | 1 (0.01%) | 522 (< 0.02%) | | 131 (< 0.02%) | 1,598 (2.27%)* | | 10 (0.17%) |
| Transcription | K | 2,405 (5.30%) | 551 (4.97%) | 140,537 (4.55%)* | | 33,959 (4.42%) | 2,992 (4.26%) | | 256 (4.3%) |
| Replication, recombination and repair | L | 4,075 (8.99%)* | 864 (7.79%) | 283,395 (9.17%)† | | 72,113 (9.39%) | 2,834 (4.03%)† | | 712 (11.95%) |
| Chromatin structure and dynamics | B | 4 (< 0.01%) | 2 (0.02%) | 764 (0.02%)* | | 132 (0.02%) | 1,026 (1.46%)* | | 17 (0.29%) |
| **Cellular processes and signaling** | | | | | | | | | |
| Cell cycle control, cell division, chromosome partitioning | D | 276 (0.61%) | 76 (0.69%) | 38,326 (1.25%)* | | 7379 (0.96%) | 1,922 (2.73%)* | | 74 (1.24%) |
| Nuclear structure | Y | 17 (0.04%) | 2 (0.02%) | 484 (0.02%) | | 121 (0.02%) | 143 (0.2%)* | | 6 (0.1%) |
| Defense mechanisms | V | 1,289 (2.84%)* | 205 (1.85%) | 99,417 (3.22%)* | | 23,244 (3.03%) | 572 (0.81%)† | | 131 (2.20%) |
| Signal transduction mechanisms | T | 497 (1.10%) | 116 (1.05%) | 127,352 (4.12%) | | 31,408 (4.09%) | 12,149 (17.28%)* | | 209 (3.51%) |
| Cell wall/membrane/envelope biogenesis | M | 3,024 (6.67%)* | 443 (4%) | 272,998 (8.83%)* | | 60,401 (7.86%) | 1,444 (2.05%)† | | 368 (6.17%) |
| Cell motility | N | 203 (0.45%) | 58 (0.52%) | 23,119 (0.75%)* | | 5,023 (0.65%) | 93 (0.13%)† | | 53 (0.89%) |
| Cytoskeleton | Z | 15 (0.03%) | 7 (0.06%) | 1,889 (0.06%) | | 437 (0.06%) | 2,647 (3.77%)* | | 102 (1.71%) |
| Extracellular structures | W | 14 (0.03%) | 6 (0.03%) | 1,259 (0.04%) | | 299 (0.04%) | 95 (0.14%)* | | 3 (0.05%) |
| Intracellular trafficking, secretion, and vesicular transport | U | 365 (0.81%) | 93 (0.81%) | 49,212 (1.59%)* | | 10,953 (1.43%) | 5,340 (7.6%)* | | 164 (2.75%) |
| Posttranslational modification, protein turnover, chaperones | O | 2,054 (4.53%) | 465 (4.53%) | 123,184 (3.99%)* | | 29,880 (3.89%) | 8,605 (12.24%)* | | 264 (4.43%) |
| **Metabolism** | | | | | | | | | |
| Energy production and conversion | C | 3,568 (7.87%)† | 1,060 (9.56%) | 132,233 (4.28%)† | | 35,679 (4.65%) | 1,440 (2.05%)† | | 379 (6.36%) |
| Carbohydrate transport and metabolism | G | 1,449 (3.20%) | 370 (3.34%) | 216,126 (6.99%) | | 61,013 (7.94%) | 2,025 (2.88%)† | | 283 (4.75%) |
| Amino acid transport and metabolism | E | 3,933 (8.68%)† | 1,052 (9.56%) | 190,552 (6.17%)† | | 52,364 (6.82%) | 1,293 (1.84%)† | | 377 (6.33%) |
| Nucleotide transport and metabolism | F | 1,582 (3.49%)† | 440 (3.97%) | 96,570 (3.13%)* | | 22,790 (2.97%) | 650 (0.92%)† | | 179 (3%) |
| Coenzyme transport and metabolism | H | 3,135 (6.92%)† | 853 (7.69%) | 77,096 (2.49%)† | | 20,123 (2.62%) | 390 (0.55%)† | | 176 (2.95%) |
| Lipid transport and metabolism | I | 913 (2.01%) | 200 (1.80%) | 64,209 (2.08%) | | 16,053 (2.09%) | 1,732 (2.46%) | | 131 (2.2%) |
| Inorganic ion transport and metabolism | P | 2,334 (5.15%) | 583 (5.26%) | 123,568 (4%)† | | 35,631 (4.64%) | 1,565 (2.23%)† | | 276 (4.63%) |
| Secondary metabolites biosynthesis, transport and catabolism | Q | 502 (1.11%) | 109 (0.98%) | 26,161 (0.85%)† | | 7,000 (0.91%) | 1,610 (2.29%)* | | 68 (1.14%) |
| **Poorly characterized** | | | | | | | | | |
| Function unknown | S | 10,164 (22.42%) | 2,407 (21.71%) | 781,616 (25.29%)† | | 195,710 (25.48%) | 13,399 (19.06%) | | 1,107 (18.57%) |
| **Total COG/KOG Annotated Genes** |  | **45,336** | **11,086** | **3,090,096** | | **768,114** | **70,289** | | **5,960** |

* Over-represented genes, relative to the long-read dataset (Fisher’s Exact Test, *P* < 0.05)

† Under-represented genes, relative to the long-read dataset (Fisher’s Exact Test, *P* < 0.05)

Table S12 (Viral-host contig associations):

Table S13 (ARG allele detections):

Table S14 – Data sequenced for this survey

| Platform | Number of flowcells | Total cumulative bases |
| --- | --- | --- |
| Illumina (short-read) NextSeq 500 | 1 (four lanes) | 172 gigabases |
| PacBio RSII | 8 | 7.57 gigabases |
| PacBio Sequel | 21 | 45.35 gigabases |

Table S15 – Rumen short-read WGS datasets used in binning or comparisons in this study.

| SRA Accession | Total reads | Short-read mapping (%) | Long-read mapping (%) | Used in MetaBat binning? | Used in Hypergeometric enrichment test?^1^ |
| --- | --- | --- | --- | --- | --- |
| PRJEB10338 | 832,702,032 | 62 | 38 | Yes | Yes |
| PRJEB21624 | 5,756,832,251 | 70 | 47 | Yes | Yes |
| PRJEB8939 | 329,335,090 | 71 | 48 | Yes | Yes |
| PRJNA214227 | 1,571,755,419 | 61 | 41 | Yes | Yes |
| PRJNA291523 | 281,175,435 | 79 | 65 | Yes | Yes |
| PRJNA60251 | 2,647,283,000 | 36 | 20 | Yes | Yes |
| PRJNA255688 | 16,506,517 | 97 | 79 | Yes | No |
| PRJNA270714 | 67,813,340 | 66 | 35 | Yes | No |
| PRJNA280381 | 95,781 | 99 | 99 | Yes | No |
| PRJNA366460 | 44,383,305 | 82 | 57 | Yes | No |
| PRJNA366463 | 22,354,786 | 84 | 65 | Yes | No |
| PRJNA366471 | 23,525,510 | 82 | 63 | Yes | No |
| PRJNA366487 | 21,116,112 | 87 | 57 | Yes | No |
| PRJNA366591 | 45,294,312 | 81 | 63 | Yes | No |
| PRJNA366667 | 20,775,651 | 90 | 68 | Yes | No |
| PRJNA366681 | 22,410,058 | 84 | 61 | Yes | No |
| PRJNA398239 | 11,758,630 | 99 | 99 | Yes | No |
| PRJNA507739 (this study) | 1,148,721,358 | 85 | 47 | Yes | Yes |

^1^ Datasets that were comprised of over 100 million WGS short-reads were used in the enrichment test.

**Figure S1 – PacBio read alignments to Long-read assembly partitioned by GC%**

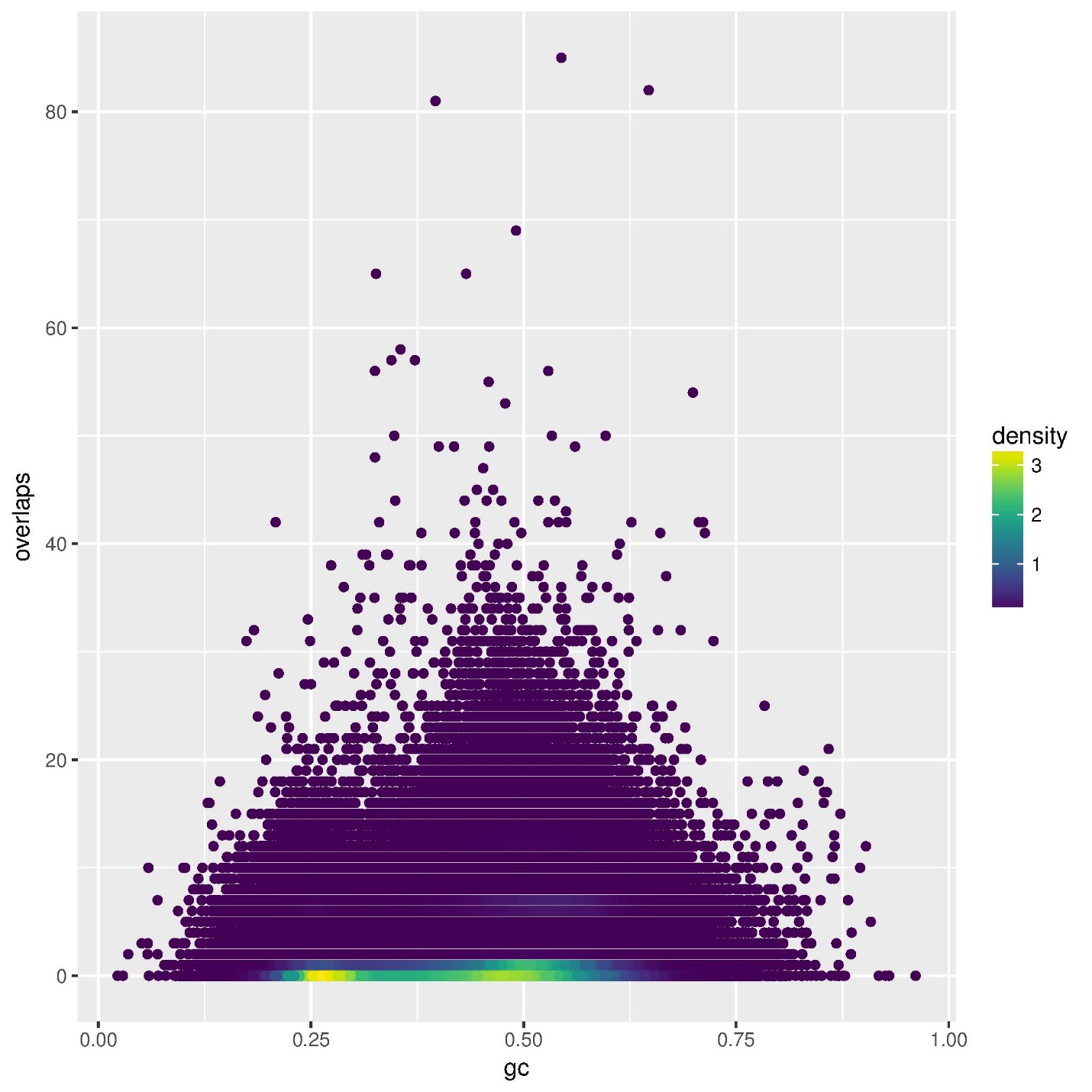

**Figure S2 – Short-Read assembly contig length vs GC% confidence interval plot**

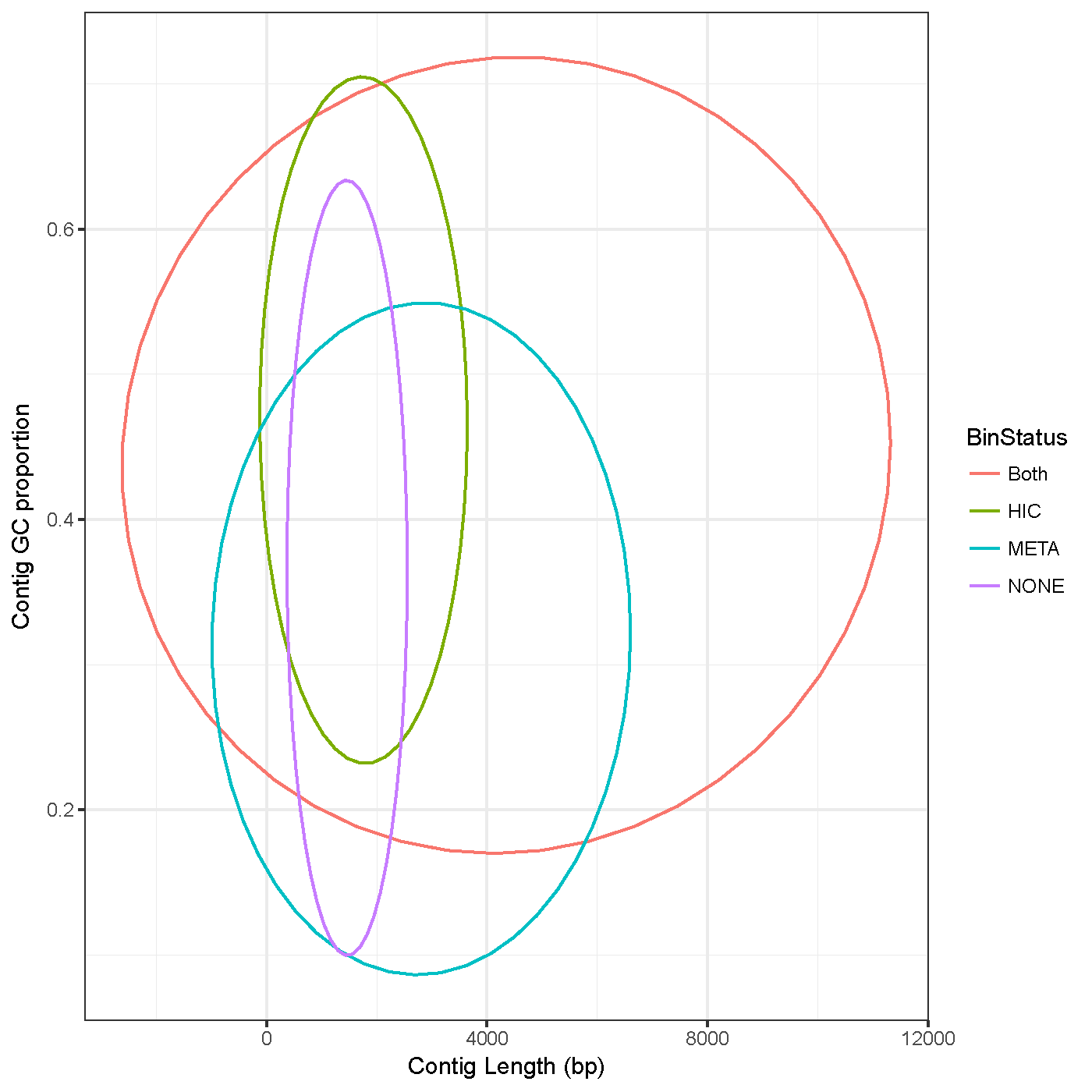

Figure S3 – Krona plot of short-read assembly contig taxonomic assignment

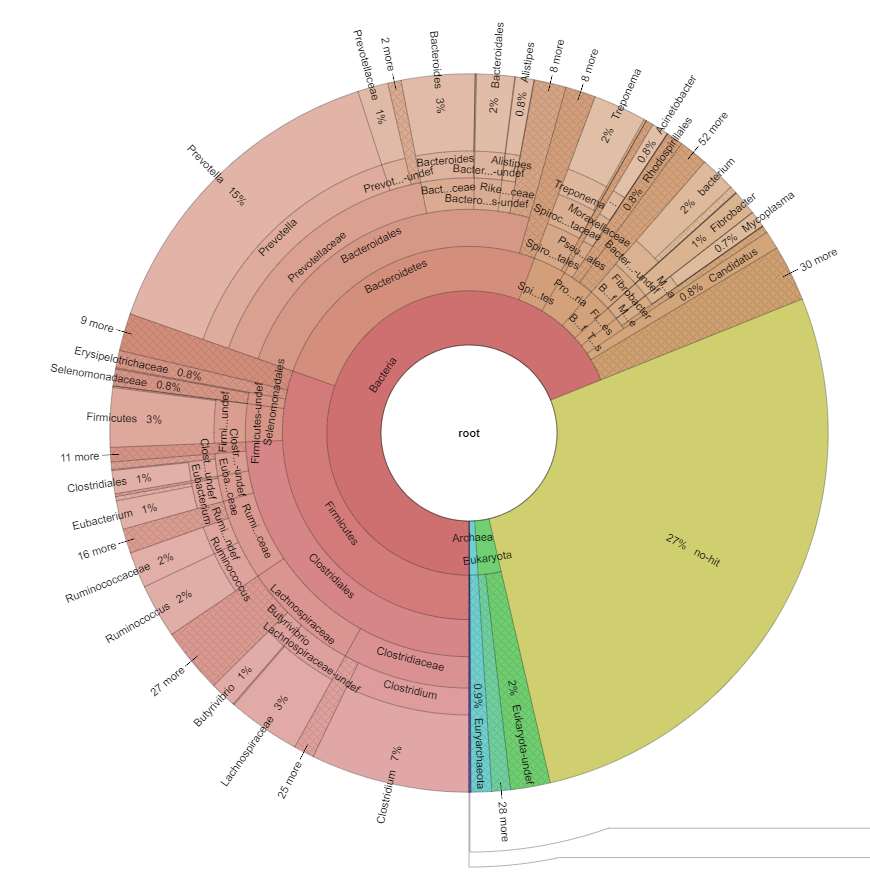

Figure S4 – Krona plot of long-read assembly contig taxonomic assignment

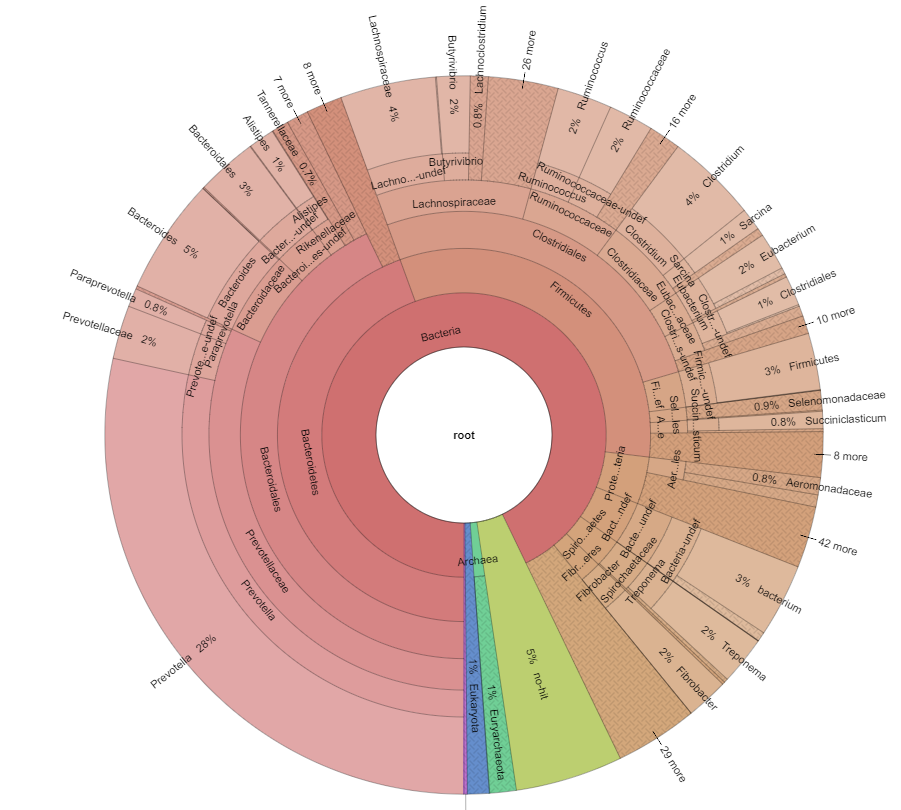

Figure S5 – Proportional COG category comparisons between short- and long-read assemblies. The 95% confidence intervals are delimited by the gray background in each plot.

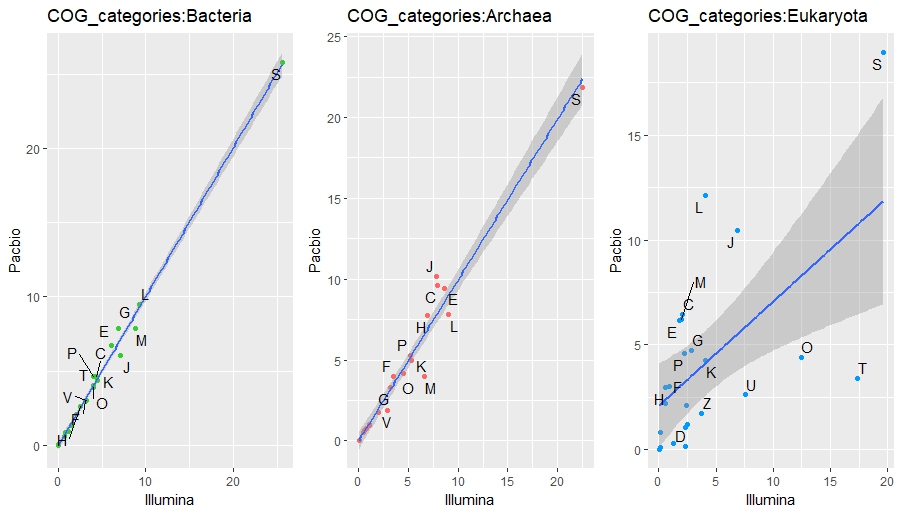

Figure S6 – Hi-C intercontig link association of ARG allele-containing and other contigs

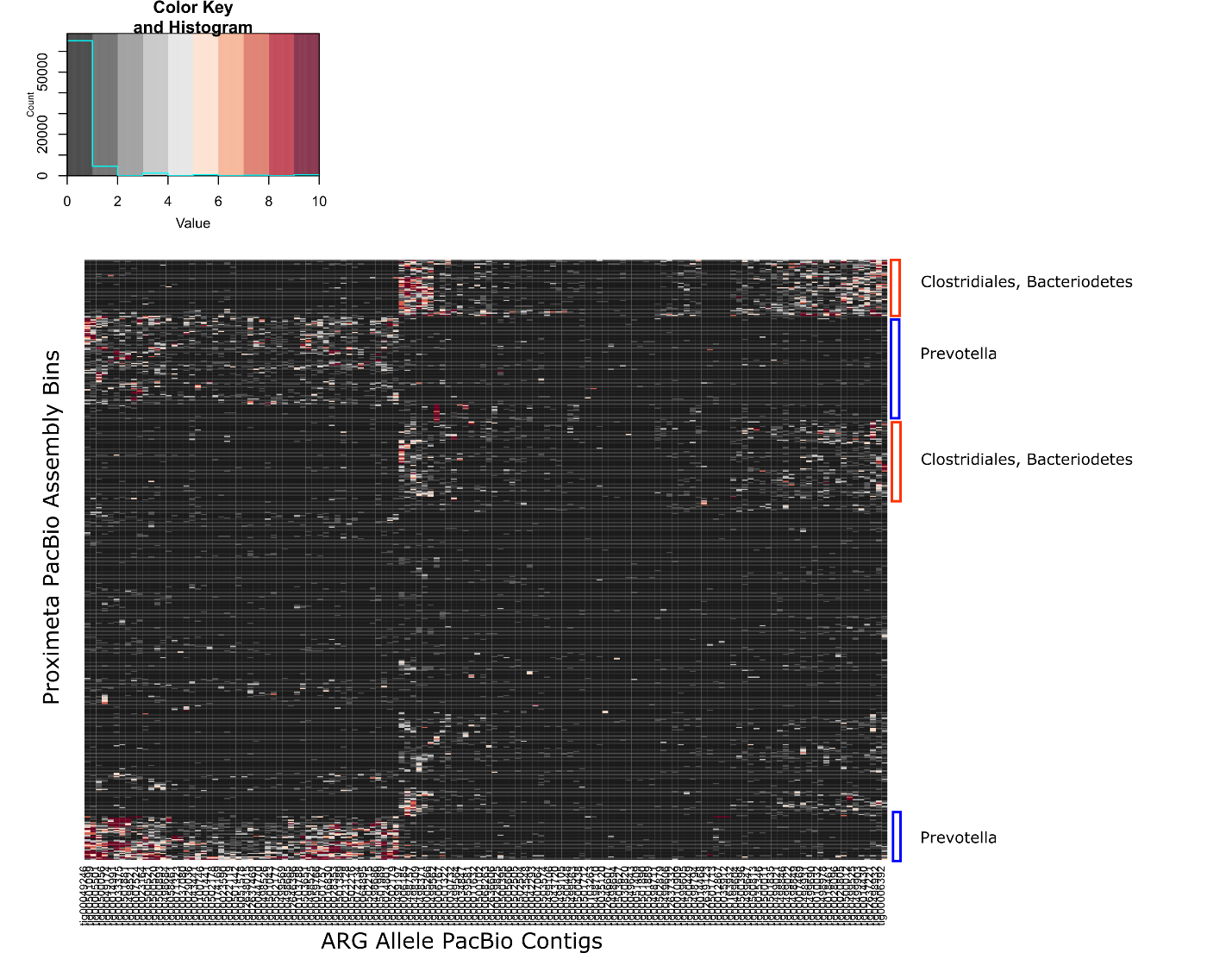

Figure S8 – Fragment analyzer spectra for rumen contents sample prepared via modified Yu and Morrison protocol.

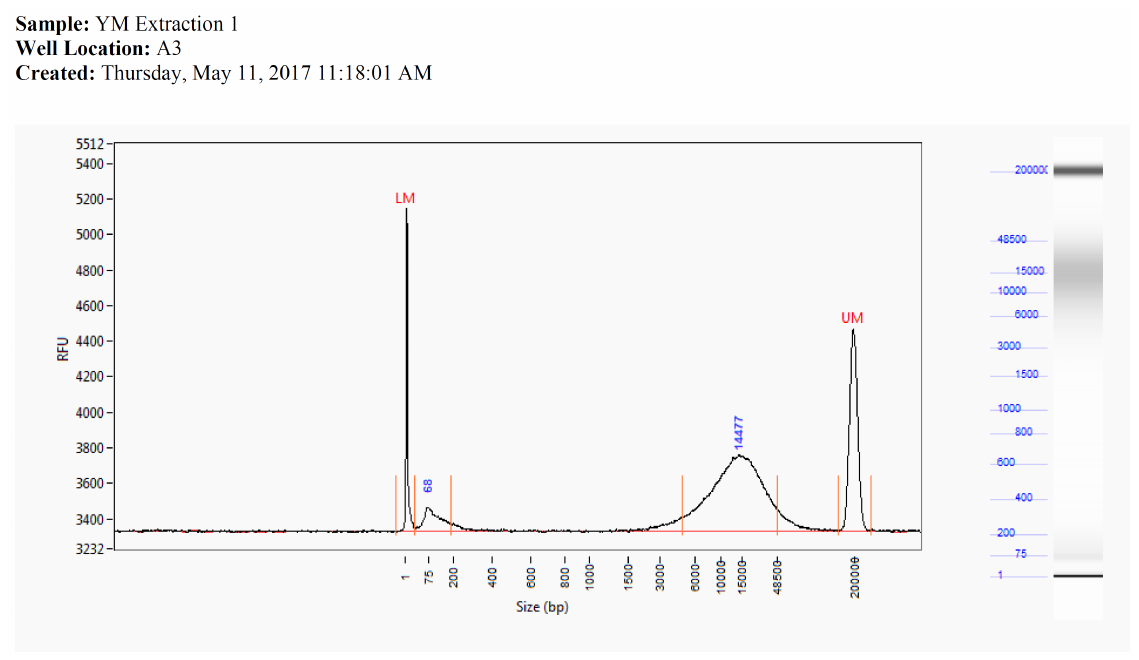

Figure S9 – short-read assembly CheckM bin statistics

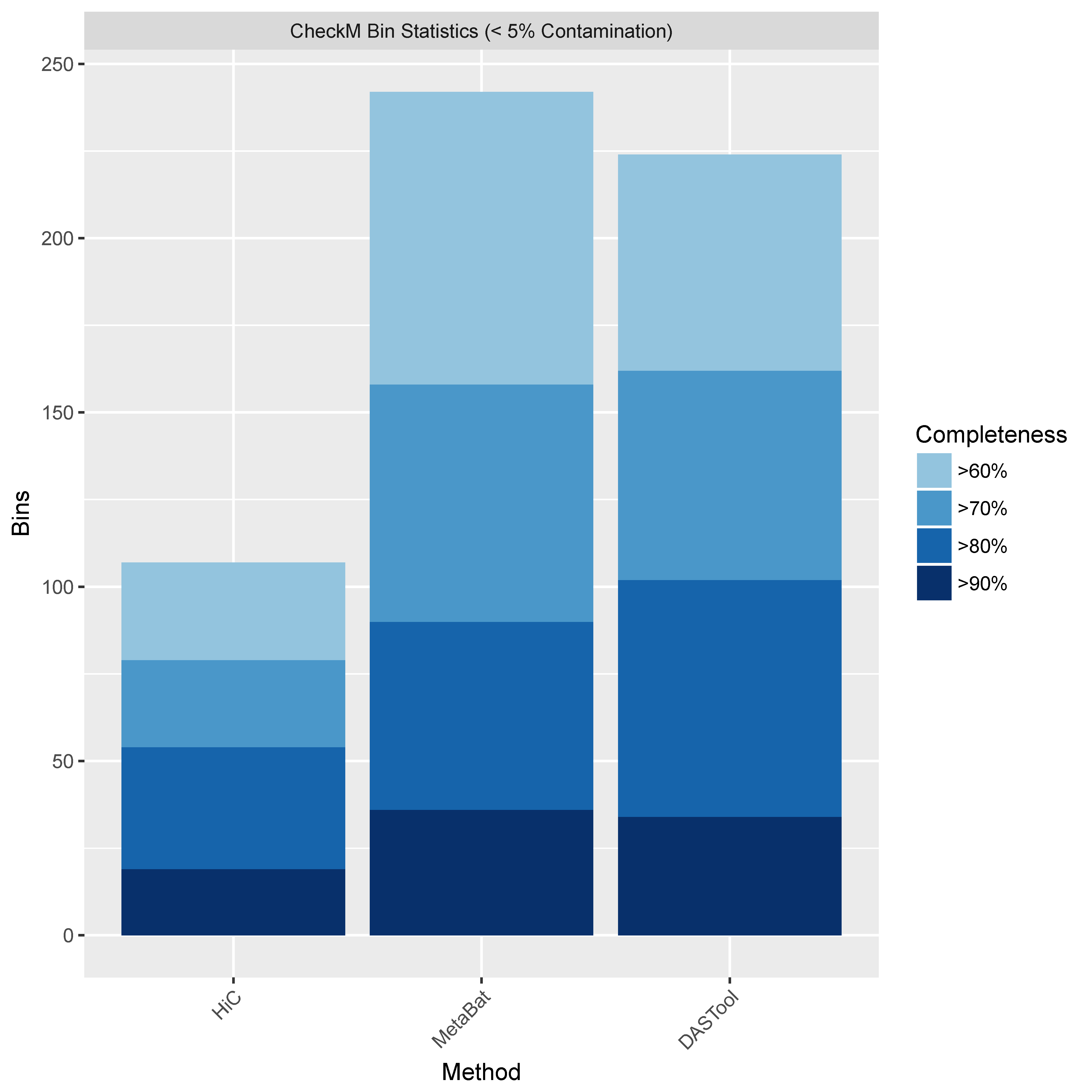

Figure S10 – Long-read assembly CheckM bin statistics

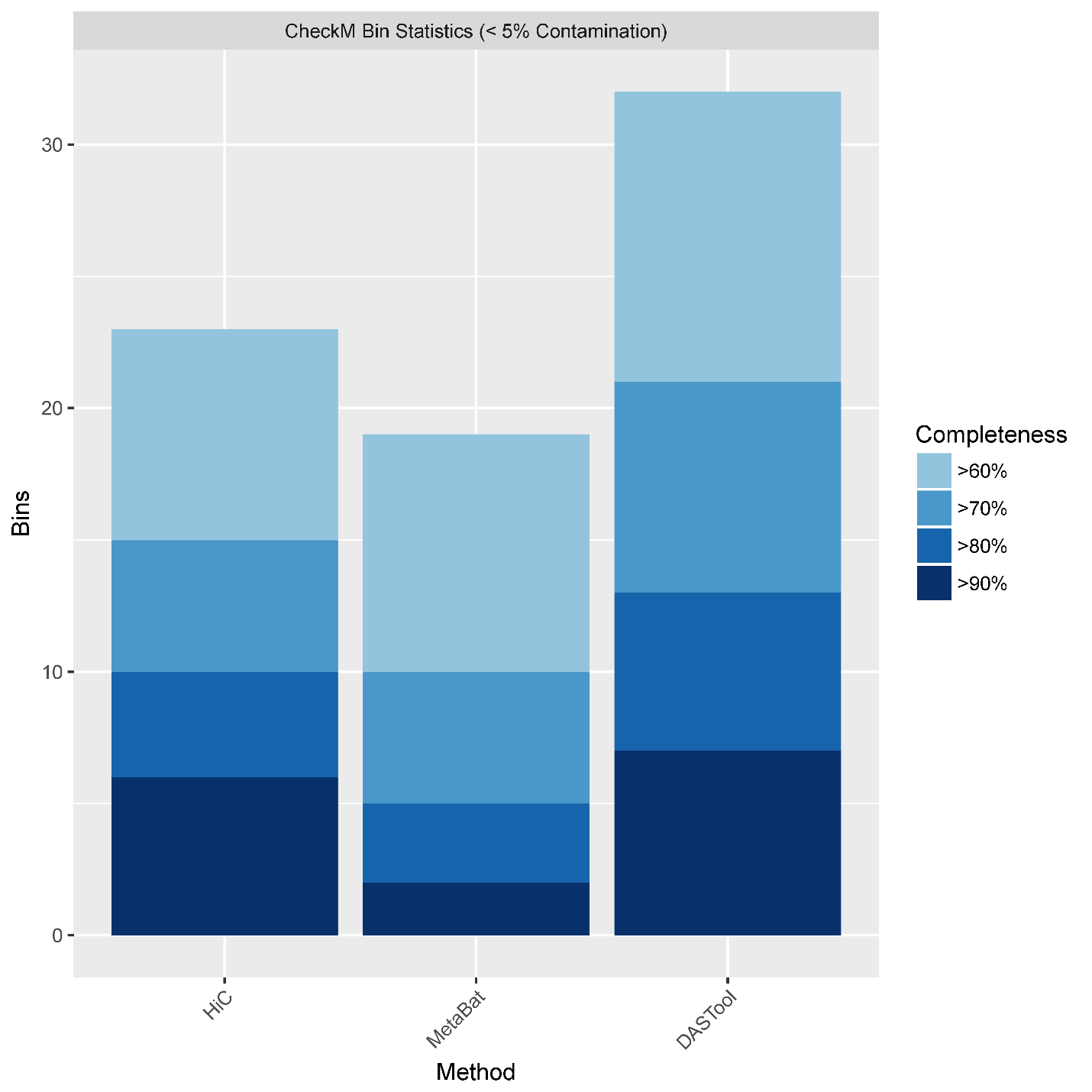

**Supplementary methods**

**Binning strategies**

In order to associate disparate contigs into biologically relevant bins, we used one computational and one technical method. We used MetaBat2(1)(referred to as “MetaBat” hereafter) to group contigs by tetranucleotide frequency and sequence coverage metrics. Sequence coverage was generated for each assembly by aligning paired-end short reads from our input dataset and 16 public SRA datasets (Table S14) using BWA MEM(2) and Samtools(3). Coverage statistics were generated from each alignment BAM file via the use of the jgi_summarize_bam_contig_depths executable provided in the MetaBat2 distribution, using default settings and the “—outputDepth” argument. MetaBat2 was run with the combined coverage statistics data on the short-read and long-read assemblies, respectively. Contig-bin assignments were tabulated from the resulting MetaBat fasta files using this custom script:

perl -e '@f = `ls public_metabat/*.fa`; chomp(@f); foreach $h (@f){@hsegs = split(/\./, $h); open(IN, "< $h"); while(<IN>){if($_ =~ /^>/){chomp; $_ =~ s/>//; print "$_\t$hsegs[-2]\n";}} close IN;}' > bin_ids.tab

Hi-C bins were generated by Phase Genomics (Seattle, WA) using their ProxiMeta analysis service. Briefly, this technique uses proximity ligation (Hi-C) connectivity between contigs to group contigs into bins. For more details about ProxiMeta clustering and analysis see (4).

Finally, we used the DAS_Tool pipeline(5) to consolidate the MetaBat and Hi-C bins and to assess bin quality using single-copy-gene (SCG) annotations. DAS_Tool was run with Diamond(6) as the default search engine and used the same Prodigal(7) ORF predictions identified through the BlobTools(8) pipeline to identify SCGs. We also ran DAS_Tool without a minimum reporting bin_score threshold so that statistics could be generated on each consolidated binset:

DAS_Tool -i pacbio_final_public_hic.unsorted.bins,pacbio_final_public_metabat.unsorted.bins -c usda_pacbio_second_pilon_indelsonly.fa -o pacbio_final_dastool -l HiC,metabat --search_engine diamond -t 10 --db_directory /mnt/nfs/nfs2/bickhart-users/binaries/DAS_Tool/db --write_bins 1 --proteins pacbio_final_prodigal_proteins.faa --score_threshold 0

DAS_Tool -i illumina_megahit_hic.unsorted.bins,illumina_megahit_public_metabat.unsorted.bins -c mick_megahit_final_full.rfmt.fa -o illumina_megahit_dastool -l HiC,metabat --search_engine diamond -t 10 --db_directory /mnt/nfs/nfs2/bickhart-users/binaries/DAS_Tool/db --write_bins 1 --proteins illumina_megahit_prodigal_proteins.faa --score_threshold 0

**Hybrid genome assembly**

As we had substantially greater depth of coverage with the short-read dataset, we attempted to scaffold our short-read assembly contigs with our long-read data using the Opera-LG scaffolder(9). Due to the constraints placed on Opera-LG by its optimized data structures, we were unable to use it to scaffold the entire short-read assembly as that assembly was larger than the 4 gigabase limit on input assemblies. To circumvent this limitation, we removed all contigs less than 1.5 kb from the short-read assembly before scaffolding. We observed only a minor improvement in scaffold N50 length and a substantial increase in N-base gap content in the hybrid assembly. Despite the longer read lengths in our long-read dataset, there was insufficient overlap and coverage on the short-read assembly contigs to generate substantially longer scaffolds.

**PacBio alignment and Hi-C inter-cluster link analysis for Viral host-specificity analysis**

Error corrected read alignments of PacBio reads were also used to identify and confirm viral host-specificity in assembled contigs from both assemblies. Filtered alignments of long-reads that spanned BlobTools/DIAMOND designated viral contigs and non-viral contigs resulted in a list of 64 and 45 candidate insertions of long-read and short-read viral contigs, respectively, in other non-viral contigs. In order to identify these associations, we first filtered read alignments of error-corrected PacBio reads to an exclusive reference set of viral contigs and identified the segments of the reads that were overhanging the end of the alignment:

minimap2 -x map-pb pacbio_pilon_viruses.fa rumen_pacbio_corrected.fasta.gz > pacbio_pilon_viruses_ecpbreads.paf

perl selectLikelyViralOverhangs.pl pacbio_pilon_viruses_ecpbreads.paf pacbio_pilon_viruses_ecpbreads.filt

We then kept all read overhangs that were greater than 150 bp in length, and extracted the portion of the error corrected reads that were previously unmapped to the viral contigs:

perl -lane 'if($F[2] - $F[1] > 150){print $_;}' < pacbio_pilon_viruses_ecpbreads.filt.subread.bed > pacbio_pilon_viruses_ecpbreads.filt.subread.gt150.bed

perl -e '@list; while(<>){chomp; @s = split(/\t/); push(@list, "$s[0]:$s[1]-$s[2]"); if(scalar(@list) > 500){print "Printing...\n"; system("samtools faidx ../../sequence_data/pilot_project/pacbio/rumen_pacbio_corrected.fasta " . join(" ", @list) . " >> pacbio_pilon_viruses_ecpbreads.filt.subread.gt150.fa"); @list = ();}} system("samtools faidx ../../sequence_data/pilot_project/pacbio/rumen_pacbio_corrected.fasta " . join(" ", @list) . " >> pacbio_pilon_viruses_ecpbreads.filt.subread.gt150.fa");' < pacbio_pilon_viruses_ecpbreads.filt.subread.gt150.bed

The overhanging read data was realigned to the entire assembly and Blobtools-derived taxonomic information was used to filter and label the association data.

perl -e 'chomp(@ARGV); open(IN, "< $ARGV[0]"); %data; while(<IN>){chomp; @s = split(/\t/); if($s[11] == 0){next;} $s[0] =~ s/\:\d+\-\d+$//; push(@{$data{$s[0]}}, $s[5]);} close IN; open(IN, "< $ARGV[1]"); while(<IN>){chomp; @s = split(/\t/); if(exists($data{$s[0]})){push(@{$data{$s[0]}}, $s[5]);}} close IN; foreach my $k (keys(%data)){print "$k\t" . join("\t", @{$data{$k}}) . "\n";}' pacbio_pilon_viruses_ecpbreads.filt.subread.gt150.paf pacbio_pilon_viruses_ecpbreads.paf > pacbio_pilon_viruses_ecpbreads.assoc.filt.tab

perl generateViralAssociationGraph.pl pacbio_secpilon_blobplot_all.pacbio_secpilon_blobplot.blobDB.table.txt pacbio_pilon_viruses_ecpbreads.assoc.filt.stringent.tab pacbio_pilon_viruses_ecpbreads.assoc.filt.stringent.cyto.tab

In order to identify Hi-C inter-cluster links, we first generated a bipartite inter-contig Hi-C read graph by aligning Hi-C read pairs to each assembly and by selecting alignments where one read pair aligned to a viral contig and the other aligned to a non-viral contig. We then created a sum value for the cardinal counts of each unique association, and filtered the resulting data by the magnitude of the count. We chose values of 10 and 20 observations as filter cutoffs for the short-read and long-read viral-host graphs, respectively, as those values represented the median counts for each dataset. This filtered data was used to generate an extended table file similar to the PacBio read alignment data listed above:

perl generateViralAssociationGraph.test.pl ../blobtools/illumina_blobplot_all.illumina_megahit_blobplot.blobDB.table.txt illumina_viruses.hiclinks.filt.tab illumina_viruses.hiclinks.filt.cyto.tab

Finally, the Hi-C inter-cluster links and PacBio read alignments were combined into a final association file using a simple overlap of the viral-host associations discovered in each separate analysis.

perl combineViralSignal.pl pacbio_pilon_viruses.hiclinks.filt.cyto.tab pacbio_pilon_viruses_ecpbreads.stringent.cyto.tab > pacbio_pilon_viruses.combined.cyto.tab

The resulting table was loaded into Cytoscape version 3.6.1(10), with the viral contigs and non-viral contigs selected as nodes, and the method of detection {“PacBio”, “HIC”, “BOTH”} selected as the edge/interaction. In the case of the long-read assembly data, viral nodes that had more than two connections with non-viral nodes were grouped using an “Attribute circle” layout. The short-read assembly had fewer of these high connectivity nodes, so the default network layout was used to display the network.

**Hypergeometric determination of contig coverage enrichment**

Short-read sequence data alignments from six of the 16 SRA datasets used to bin the contigs (see Table S14) were used in the subsequent analysis of coverage enrichment. We calculated read coverage enrichment only for the AN bins in the short- and long-read assemblies in the following analysis. In each test, we calculated the proportion of the total aligned reads from each respective dataset to each contig from the short-read and long-read AN bins. We grouped contigs by their likely genus origin based on their Blobtools taxonomic designation. We then identified cases where our short-read dataset had the most or the least aligned reads to each contig and classified them as “max” or “min” events, respectively. Grouping contigs by their genus designation, we then tested whether the proportion of “max” or “min” read depth contigs observed were higher than expected in the group using the Scipy hypergeometric survivor function (scipy.stats.hypergeom.sf). To correct for multiple hypothesis testing, we applied a Benjamini-Hochberg correction to the hypergeometric survivor function p values using an alpha of 0.05.
